# BRD4 binds the nucleosome via both histone and DNA interactions

**DOI:** 10.1101/2025.05.29.656846

**Authors:** Jiang Zhu, Erik M. Leith, Erin N. O’Donnell, Bryan P. Manzano, Shwu-Yuan Wu, Cheng-Ming Chiang, Jean-Paul Armache, Song Tan

**Affiliations:** Center for Eukaryotic Gene Regulation, Department of Biochemistry and Molecular Biology, The Pennsylvania State University, University Park, PA 16802, USA; Simmons Comprehensive Cancer Center, University of Texas Southwestern Medical Center, Dallas, TX 75390, USA; Department of Biochemistry, University of Texas Southwestern Medical Center, Dallas, TX 75390, USA; Department of Pharmacology, University of Texas Southwestern Medical Center, Dallas, TX 75390, USA

**Author notes:** Contributed equally to this work (Jiang Zhu, Erik M. Leith).

## Abstract

BRD4, a bromodomain and extraterminal (BET) family transcriptional regulator of cell cycle progression, cell differentiation and cancer development, is believed to be recruited to chromatin via interactions between its tandem bromodomains (BD1 and BD2) and acetylated histone tails. Although extensive studies have explained how individual BRD4 bromodomains bind to acetylated peptides and how BET inhibitors interfere with such interactions, equivalent studies of full-length BRD4 protein with the nucleosome have been lacking. Our cryo-EM structure of the BRD4 short (BRD4-S) isoform bound to a nucleosome diacetylated on histone H4 shows how BRD4 BD1 engages both the H4 tail and nucleosomal DNA. Unexpectedly, our biochemical studies indicate that BRD4 uses basic regions outside of the bromodomains to bind nucleosomes tightly even in the absence of histone acetylation. Our results further show that histone H4 acetylation influences the conformation of the BRD4/nucleosome complex.

## Introduction

BRD4, a member of the bromodomain and extraterminal (BET) family of gene regulatory proteins, has fundamental roles in transcriptional regulation, DNA replication, DNA repair, cell cycle progression, cell differentiation and cancer development^1–6^. Like other BET family members BRD2, BRD3 and BRDT, it contains tandem bromodomains (BD1 and BD2), which bind acetylated lysine residues of target proteins such as histones^7–11^, and the extraterminal (ET) domain which recruits other chromatin effector proteins^12–15^. BRD4 is usually thought to recruit the positive transcriptional elongation factor b (P-TEFb) kinase complex which enables release of RNA polymerase II (Pol II) from a transcriptional paused state to a transcriptional elongation state^16–18^. However, recent data indicate that BRD4 may not be required for P-TEFb localization^19–22^ and that BRD4 may function instead to assemble functional Pol II elongation complexes^19,21,23^.

Three isoforms of BRD4, two short (S) and one long (L), are present in human cells: BRD4-S(a), BRD4-S(b) and BRD4-L^24,25^. All three contain the same N-terminal 719 residues including the BD1, BD2 and ET domains. BRD4-S(a) (hereafter abbreviated as BRD4-S) and BRD4-L exhibit opposing functions in breast cancer, with BRD4-S acting as an oncogene and BRD4-L as a tumor suppressor^25^. All three BRD4 isoforms are believed to be targeted to acetylated chromatin through interactions between BD1/BD2 and acetylated histones^1,25,26^, but BRD4-S has been found to bind more tightly to acetylated histone H4 than BRD4-L^27^. Extensive structural and biochemical studies have characterized how bromodomains use a hydrophobic cavity to bind acetyl-lysine peptides and how BET inhibitors such as JQ1 compete for binding to this hydrophobic cavity^28–35^. BRD4’s role in oncogenesis and its overexpression in cancer cells make it an anti-cancer target, and JQ1 and other BET inhibitors have been shown to slow tumor growth^2,31,36,37^.

The vast majority of the structural and biochemical studies between BRD4 and chromatin have focused on the interactions between individual BRD4 bromodomains and histone peptides. To address the question of how the BRD4 protein interacts with acetylated histones in the context of the nucleosome and not just histone peptides, we have performed structural studies of the BRD4-S protein in complex with acetylated nucleosomes. Our cryo-EM structure of the BRD4/nucleosome complex shows how BRD4 BD1 interacts with not only the histone H4 N-terminal tail but also with nucleosomal DNA. Unexpectedly, we find that the BRD4 protein does not need histone acetylation or even the H4 tail to bind with high affinity to nucleosomes. Instead, we find that histone H4 acetylation influences the conformation of the BRD4/nucleosome complex.

## Results

### BRD4/nucleosome structure

We reconstituted BRD4-S with nucleosomes containing acetylated histones at a variety of positions on histone H3 and/or H4. We then analyzed these BRD4/nucleosome complexes using cryoelectron microscopy. For most of the BRD4/nucleosome complexes we examined, we were able to observe extra density on the histone face of the nucleosome in the resulting image reconstructions, but that density was diffuse and difficult to interpret. The exception was the BRD4/nucleosome complex containing H4 K12acK16ac and H3 K18ac for which tubular density corresponding to the bromodomain four-helix bundle was clearly visible (**Fig. 1a**). The nominal resolution of the BRD4/nucleosome cryo-EM reconstruction using 80,700 particles was 2.9 Å **(Extended Data Fig. 1, 2)**, but this reflects the much better resolution for the nucleosome component of the complex compared to the local resolution of about 4.5 Å for the BRD4 bromodomain **(Extended Data Fig. 1**). Such a difference in local resolution is often observed in nucleosome complexes presumably due to motion of the chromatin factor with respect to the nucleosome^38–41^.

**Figure 1:**
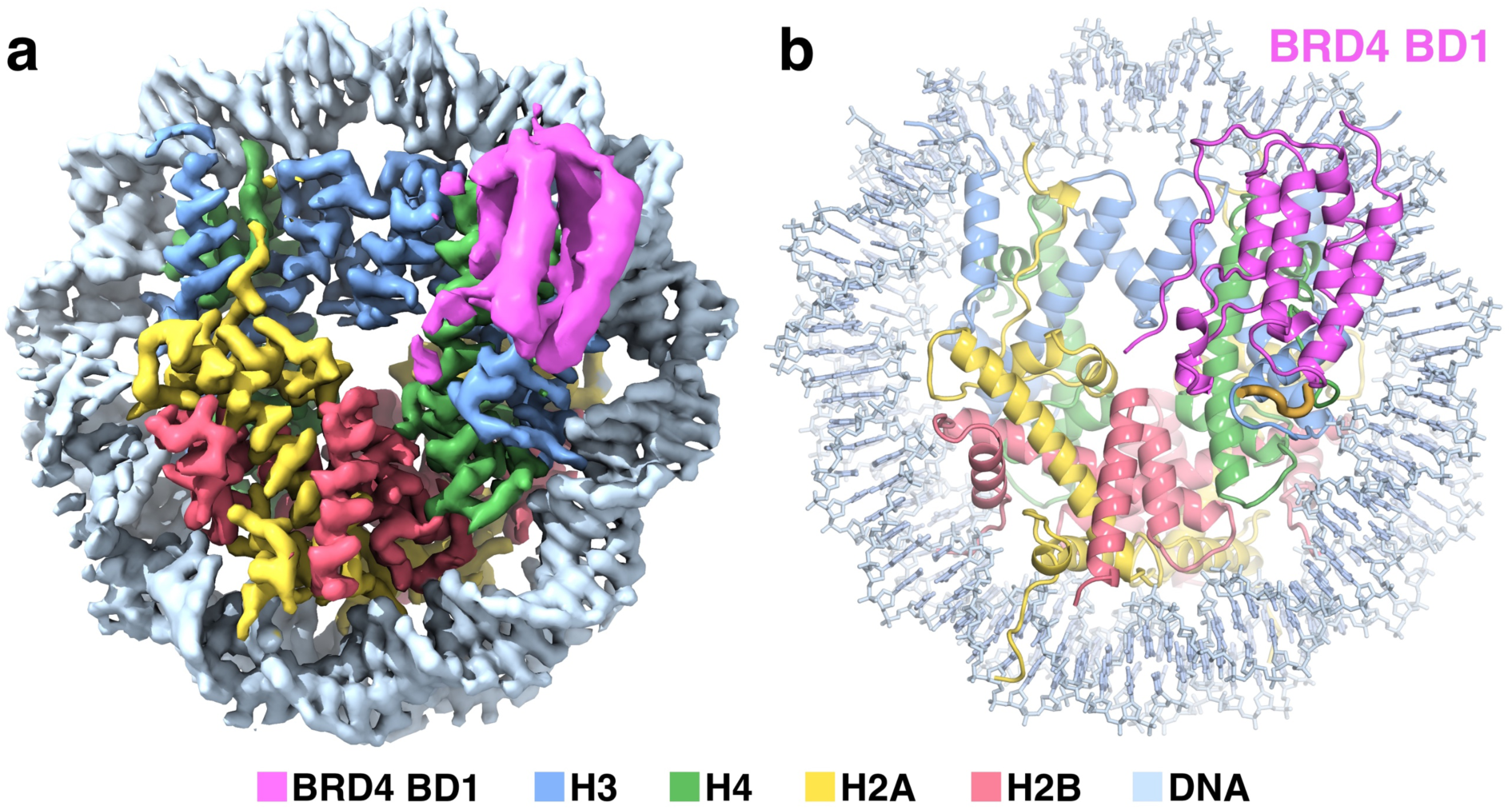
Overview of BRD4/nucleosome structure. (a) Cryo-EM map and (b) cartoon representation of the BRD4-S/nucleosome complex showing how the BRD4 BD1 (pink) interacts with the nucleosome. The modeled histone H4 tail residues 11-16 are shown as a thicker gold line.

Our structural model contains the nucleosome and the crystal structure of BRD4 BD1 built into the extranucleosomal tubular density corresponding to the bromodomain α-helices. (**Fig. 1b**). We believe the extranucleosomal tubular density corresponds to BRD4 BD1 because the structural model positions the bromodomain’s peptide binding pocket to interact with histone H4 K12acK16ac N-terminal tail, one of the preferred peptides for BRD4 BD1^42^. We used nucleosomes also containing H3 K18ac in the hope that BRD4 BD2 would engage that modified histone tail. However, we only observe density for one bromodomain in our reconstruction. This suggests the rest of the BRD4-S protein, which includes BD2 and the ET domain, is conformationally flexible on the nucleosome.

In our structural model, BRD4 BD1 is located on the histone face of the nucleosome above the histone H4 N-terminal tail. This allows the histone H4 tail to enter the BD1 peptide binding pocket, and in our cryo-EM map we observe density for the H4 tail approaching the peptide binding pocket **(Extended Data Fig. 3a**). In addition, BD1 is positioned to interact with the phosphate backbone across the minor groove of nucleosomal DNA around superhelical position (SHL) +1.5 via helix ⍺Z residues R68, K72 and K76.

### BRD4 does not require acetylated histones to bind with high affinity to nucleosomes

To further understand how the BRD4 protein interacts with the nucleosome, we performed binding studies using insights from our structural studies. To quantitatively measure how BRD4-S interacts with nucleosomes, we employed the time-resolved fluorescence resonance energy transfer (TR-FRET) nucleosome binding assay recently developed by the McGinty laboratory^43^. We initially performed these experiments at a NaCl concentration of 70 mM. Since the BRD4 bromodomains specifically bind to acetylated histone tails, we expected that BRD4-S would bind tighter to acetylated nucleosomes compared to unmodified nucleosomes. Our results show that BRD4-S binds tightly (K_d_ = 8.0 nM) to H4 K12acK16ac nucleosomes (**Fig. 2a, Extended Data Fig. 4a**). However, we find that BRD4-S binds with similar or even slightly higher affinity to unmodified nucleosomes (6.6 nM) as it does to the acetylated nucleosomes. Even more unexpected is the result that BRD4-S binds equally well to H4 tailless nucleosomes, which lack the H4 residues thought to bind to BRD4, as it does to unmodified nucleosomes (**Fig. 2a**). This indicates that BRD4-S’s affinity to the nucleosome is dictated by more than its bromodomains’ interactions with acetylated histone tails. Consistent with this conclusion is our finding that binding of individual BRD4 BD1 and BD2 bromodomains to H4 K12acK16ac nucleosomes could not be detected in our assay **(Extended Data Fig. 4a**).

**Figure 2:**
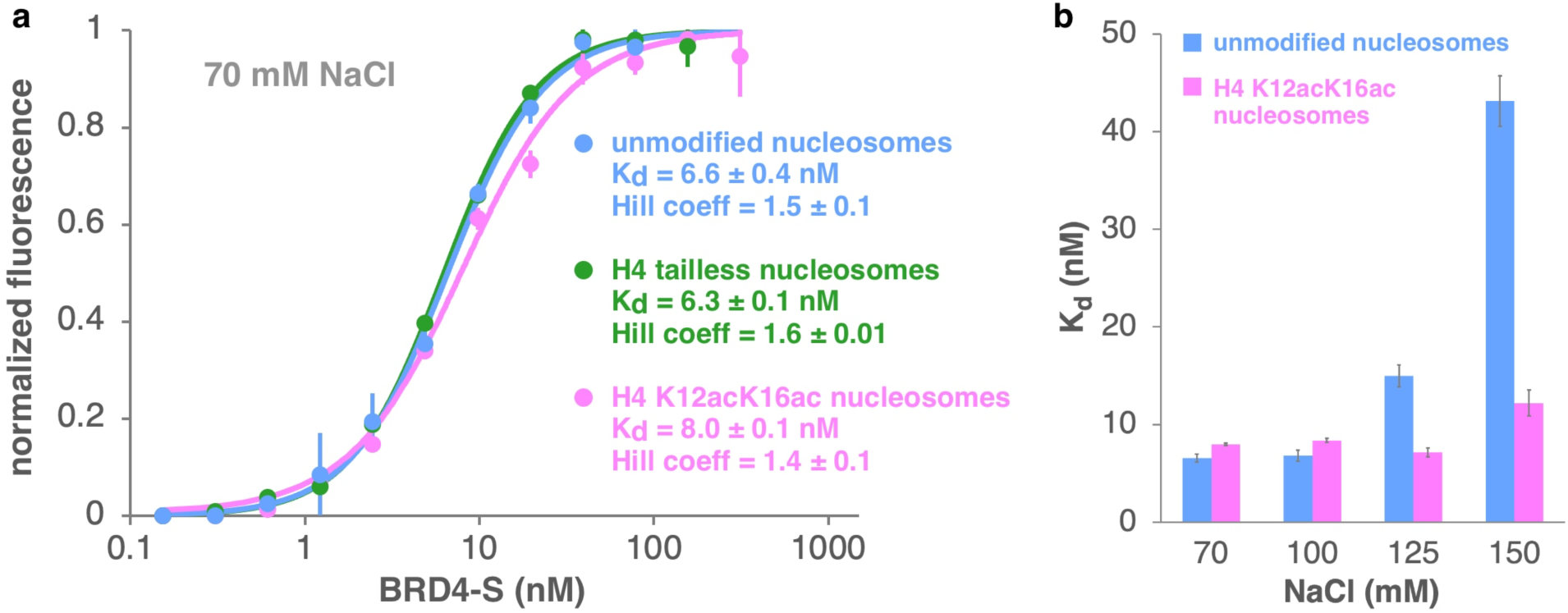
BRD4-S binds to nucleosomes with nanomolar affinity at 70 mM NaCl independent of histone acetylation. (a) Time-resolved FRET binding assay results for BRD4-S binding to unmodified (blue), H4 tailless = H4(24-102) (green) and H4 K12acK16ac nucleosomes (pink) in 70 mM NaCl. (b) Effect of NaCl concentration on BRD4-S binding to unmodified (blue) or H4 K12acK16ac nucleosomes (pink) as assayed by TR-FRET.

### Role of BRD4 basic patches in high affinity binding to nucleosomes

Given our findings that BRD4-S binds tightly to both unmodified and H4 K12acK16ac nucleosomes and that individual BRD4 bromodomains are not sufficient to bind H4 K12acK16ac nucleosomes, we asked which regions of BRD4 mediated binding to nucleosomes. When we examined the BRD4-S protein sequence, we noticed five patches enriched with Lys and Arg basic residues (**Fig. 3a**). Basic patch 1 follows almost immediately after BRD4 BD1, while basic patches 2 and 3 are positioned between basic patch 1 and BD2. Basic patch 2 is contained within the conserved BET family motif A^44^. Basic patches 4 and 5 follow the phosphorylation rich region after BD2 and are contained with the previously identified BID basic interaction domain^45^. Since a major feature of the nucleosome is the negatively charged nucleosomal DNA, we mutated the Lys and Arg residues in these BRD4-S basic patches to either Ala to remove and neutralize the basic side chain or to Glu to reverse the charge. Basic patches 4 and 5 also contain acidic residues so we only mutated the basic residues to Ala to avoid dramatically changing the charge distribution (the 20-residue basic patch 4 contains 14 basic residues and 4 acidic residues). We find that mutating individual basic patches with Ala or Glu substitutions had little effect on BRD4-S’s affinity to H4 K12acK16ac nucleosomes at 70 mM NaCl (**Fig. 3b, Extended Data Fig. 5a, Extended Data Table 3**). Mutating pairs of basic patches (2 and 3, 4 and 5) or three patches (1, 2 and 3) similarly had minor effects on nucleosome binding. However, when all five BRD4-S basic patches were mutated, we could not detect nucleosome binding. These results suggest that the five BRD4-S basic patches may have redundant roles in mediating BRD4-S’s binding to the nucleosome.

**Figure 3:**
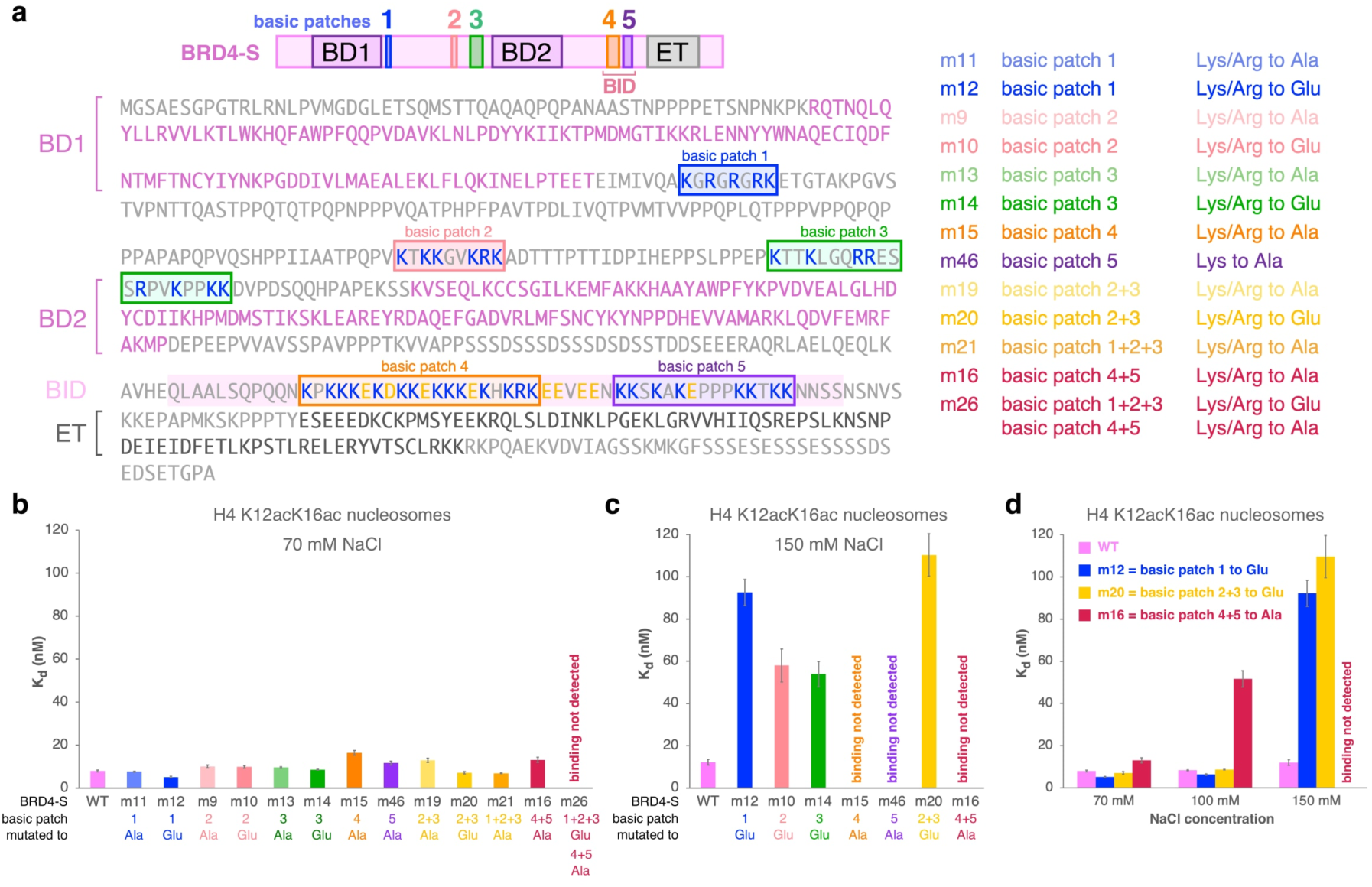
BRD4 basic patches mediate redundant interactions with H4 K12acK16ac nucleosomes. (a) BRD4-S domains and basic patches highlighted in cartoon and primary sequence (left) and identity of BRD4-S basic patch mutations studied (right), (b) TR-FRET dissociation constants for BRD4-S basic patch mutants binding to H4 K12acK16ac nucleosomes in 70 mM NaCl, (c) TR-FRET dissociation constants for BRD4-S basic patch mutants binding to H4 K12acK16ac nucleosomes in 150 mM NaCl, (d) effect of salt concentration on select BRD4-S basic patch mutations on binding to H4 K12acK16ac nucleosomes.

These results still did not explain how BRD4-S specifically recognizes acetylated nucleosomes. We reasoned that if BRD4-S uses basic residues to bind to nucleosomal DNA, these ionic interactions should depend on ionic strength. We therefore performed salt titrations to assess the ionic strength dependence of nucleosome binding by BRD4-S. Our results show that a clear dependence on ionic strength of BRD4’s affinity to H4 K12acK16ac nucleosomes compared to unmodified nucleosomes with similar affinities at 70 and 100 mM NaCl but increasing differences at 125 and 150 mM NaCl (**Fig. 2b, Extended Data Fig. 4b**). At 150 mM NaCl, BRD4-S binds to unmodified nucleosomes with a binding affinity of 43 nM versus 12 nM for H4 K12acK16ac nucleosomes for a 3.6x preference (**Extended Data Table 3**). Thus, the bromodomain-acetylated histone tail interaction in the context of the nucleosome can be detected at higher ionic strength presumably by decreasing the contributions of the charge-dependent BRD4-S basic residues-nucleosomal DNA interactions. We therefore performed subsequent nucleosome binding affinity experiments in 150 mM NaCl.

In addition to comparing binding to unmodified vs H4 K12acK16ac nucleosomes, we examined the effect of BRD4 binding at 150 mM NaCl to acetylated H4 tails in the context of the nucleosome by removing the H4 tail, by adding the JQ1 competitive inhibitor^31^ and by mutating the BRD4 bromodomain peptide binding pockets (**Fig. 4a, b, Extended Data Fig. 6, Extended Data Table 3**). We observe a similar 3- to 4-fold effect on nucleosome binding in each case. BRD4-S binds to H4 tailless nucleosomes 3.9-fold less tightly than to the H4 K12acK16ac nucleosomes. Similarly, incubating BRD4-S in the presence of JQ1, which competitively binds to the bromodomain acetylated peptide binding pocket, decreased binding affinity 3.1-fold. It is worth noting that despite this reduction in binding in the presence of the JQ1 competitive inhibitor, BRD4-S still binds acetylated nucleosomes tightly with a dissociation constant of 37 nM. The BRD4-S Y97F,N140A mutations in the BD1 acetylated peptide binding pocket, which severely diminished binding to acetyl-lysine peptides^42,46^, decreased binding to H4 K12acK16ac nucleosomes 4.1-fold. This was almost twice the effect of the equivalent BD2 mutations, which decreased binding to H4 K12acK16ac nucleosomes only 2.2-fold. As a negative control, we mutated two basic residues on BD1 facing away from the nucleosome in our cryo-EM reconstruction. This BRD4-S K99E,K102E mutant protein bound H4 K12acK16ac nucleosomes with a dissociation constant of 15.6 nM, similar to the 12.2 nM for the wild type protein. These results indicate that acetylation of histone H4 K12 and K16 contributes only modestly to the binding affinity of BRD4-S to nucleosomes.

**Figure 4:**
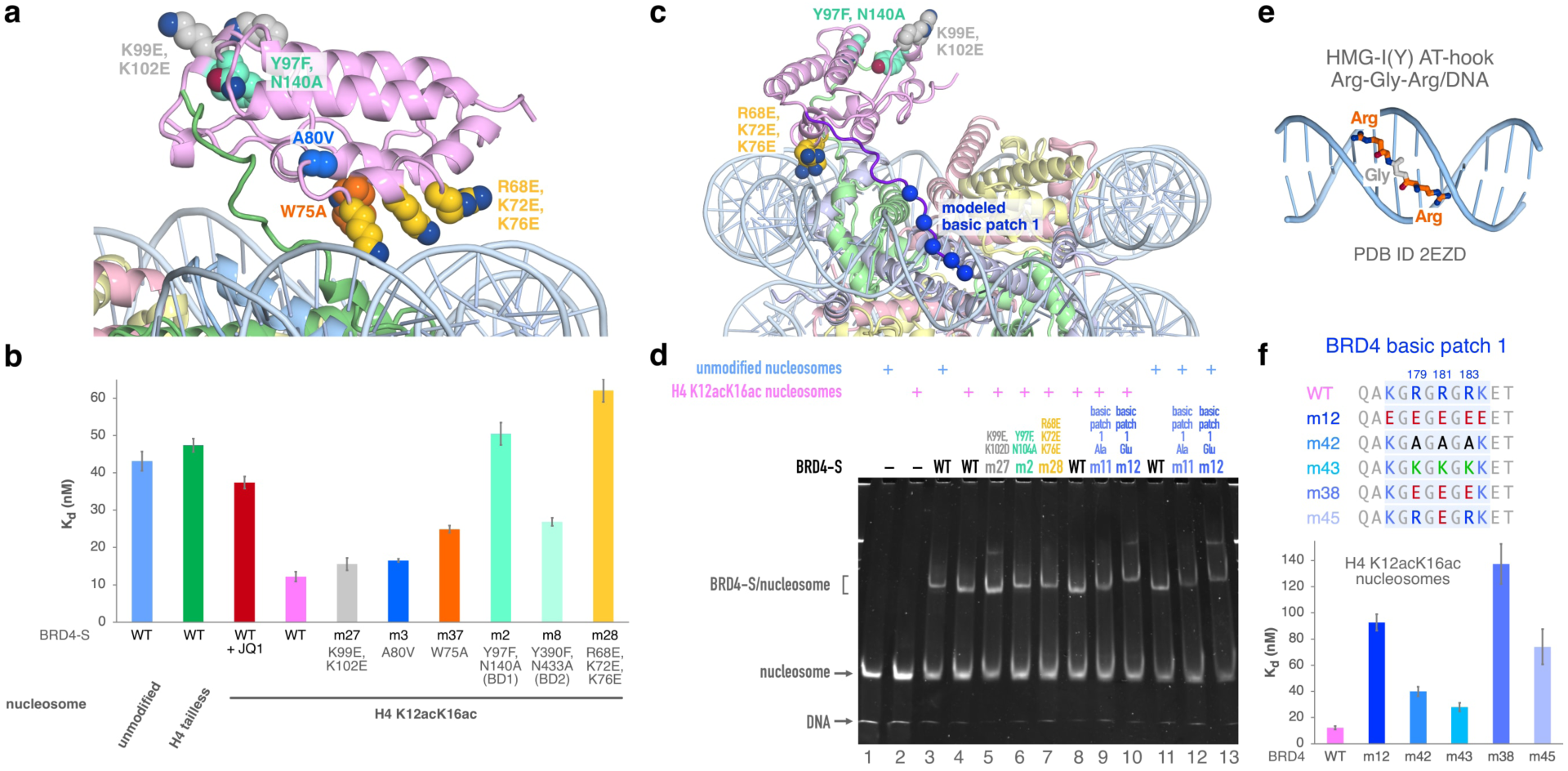
Role of BRD4 BD1 bromodomain and basic patch 1 residues in nucleosome binding. (a) BRD4 BD1 interactions with the nucleosome with key regions highlighted, (b) TR-FRET dissociation constants for BRD4-S bromodomain mutants binding to H4 K12acK16ac nucleosomes, (c) model for how BRD4 basic region 1 could interact with nucleosome DNA minor groove with C_⍺_ positions of the 5 basic residues shown in blue spheres, (d) effect of BRD4-S BD1 mutations on BRD4-S/nucleosome complex mobility in gel mobility shift assay, (e) NMR structure of HMG-I(Y) AT-hook Arg-Gly-Arg region binding to DNA (PDB ID 2EZD), protein residues outside of the Arg-Gly-Arg region not shown, (f) TR-FRET dissociation constants for BRD4-S basic patch 1 mutants binding to H4 K12acK16ac nucleosomes.

Our binding experiments also validate the interactions observed or deduced from our cryo-EM reconstruction (**Fig. 4a, b, Extended Data Fig. 6, Extended Data Table 3**). Mutating BRD4 BD1 helix ⍺Z residues R68, K72 and K76 (in position to interact with nucleosomal DNA) to Glu had a 5-fold adverse effect on BRD4-S’s binding affinity to H4 K12acK16ac nucleosomes, a greater effect than mutating the BD1 peptide binding site. The BRD4 BD1 residue W75 appears to interact with histone H4 residue R23 in our cryo-EM reconstruction of the BRD4/nucleosome complex, and we detect a two-fold decrease in binding affinity of the BRD4-S(W75A) variant to H4 K12acK16ac nucleosomes. However, another BRD4 BD1 residue, A80, which might also interact with H4 R23, showed only a 1.4-fold decrease in binding affinity, an effect similar to the K99E,K102E negative control.

We next analyzed the role of the BRD4 basic patches for binding to H4 K12acK16ac nucleosomes in 150 mM NaCl (**Fig. 3c, Extended Data Fig. 5b, Extended Data Table 3**). At this higher salt concentration, we now detect differences in binding affinity when individual BRD4 basic patches are mutated. Mutating basic patches 4 and 5 had the largest effect: binding of these mutated BRD4-S proteins to H4 K12acK16ac nucleosomes could not be detected in our TR-FRET binding assay. Partial deletion of basic patch 5 had previously been observed to adversely affect the binding of a BRD4 truncation containing BD2 and the BID to naked DNA^47^, consistent with a possible role for the BRD4 basic patch 5 to bind nucleosomal DNA. A role for basic patches 4 and 5 in nucleosome binding is also consistent with findings that BRD4(462-599) containing the BID was sufficient to bind to nucleosomes purified from a human cell line^48^. Mutating basic patch 1 residues to Glu reduced binding about 8-fold, while the equivalent mutations to basic patches 2 or 3 each reduced binding 4 to 5-fold. How much positive charges are reduced in the BRD4 mutant proteins did not always correlate with the effect on nucleosome binding. Mutating basic patch 5 yielded a BRD4-S variant with a theoretical pI of 7.56 and for which binding to acetylated nucleosomes could not be detected. In contrast, the BRD4-S protein with basic patch 3 basic residues mutated to Glu has a more acidic theoretical pI of 6.60 and this protein bound acetylated nucleosomes with a less severe 4.4-fold reduction in binding affinity. The concentration of salt affected the binding affinity of different BRD4-S basic patch mutant proteins differently. As described previously, at 70 mM NaCl all basic patch mutants examined bound with similar affinity to H4 K12acK16ac nucleosomes except when mutating all five BRD4 basic patches at the same time. At 100 mM NaCl, mutating basic patches 1 or 2+3 had very minor effects for BRD4-S binding to H4 K12acK16ac nucleosomes, whereas mutating basic patches 4+5 decreased the binding affinity 4-fold. These effects were further accentuated at 150 mM NaCl (**Fig. 3d, Extended Data Fig. 5c, Extended Data Table 3**).

We note that unlike many other nucleosome-binding proteins^49^, BRD4-S does not appear to engage the nucleosome acidic patch since mutations in the nucleosome acidic patch have no detectable effect on BRD4-S binding **(Extended Data Fig. 7**). The same mutations adversely affected binding of RCC1, a protein known to use an arginine anchor to bind to the nucleosome acidic patch^49–52^. Our results therefore suggest that BRD4-S uses its basic patches redundantly to interact with nucleosomal DNA and not with the nucleosomal histone acidic patch.

### Role of BRD4 basic patch 1 in binding nucleosomal DNA

The deleterious effect of mutating the relatively small BRD4 basic patch 1 on nucleosome binding was particularly intriguing because we observe weak density at the C-terminal end of BRD4 BD1 in a subset of particles in our cryo-EM analysis of the BRD4-S/nucleosome complex **(Extended Data Fig. 2, Extended Data Fig. 3b**). This density extends towards the nucleosomal DNA backbone around SHL+0.5 before approaching the DNA minor groove at the nucleosome dyad. We observed that the central 5 residues of the BRD4 basic patch 1 sequence (KG<u>RGRGR</u>K) contains two overlapping RGR sequences found in the AT-hook DNA-binding motif where the two Arg side chains of the RGR sequence bind across and fill the DNA minor groove (**Fig. 4e**)^53^. We have therefore modeled the BRD4 basic patch 1 in the minor groove (**Fig. 4c**). Our biochemical studies show that the three Arg residues in the BRD4 basic patch 1, R179, R181 and R183, play important roles in BRD4-S binding to the nucleosome. Mutating just these three Arg residues to Glu was sufficient to reduce binding affinity to the H4 K12acK16ac nucleosomes 11-fold (**Fig. 4f, Extended Data Fig. 6, Extended Data Table 3**). Furthermore, the basic charge of these three Arg residues is not sufficient for full binding activity since replacing these three Arg residues with Lys led to a 2.3-fold reduction in BRD4-S binding affinity to H4 K12acK16ac nucleosomes. This suggests that one or more of the three Arg side chains specifically interacts with the nucleosome, potentially in the DNA minor groove.

### The conformation of the BRD4/nucleosome complex depends on histone acetylation

In addition to using time-resolved FRET, we employed the native polyacrylamide gel electrophoresis (PAGE) gel mobility shift assay to analyze BRD4-S’s binding to the nucleosome. We find that the BRD4-S/nucleosome complex migrates slightly faster when H4 K12acK16ac nucleosomes instead of unmodified nucleosomes are used (**Fig. 4d: lanes 3 and 4)**. This faster mobility does not appear to be attributed to the acetylation charge neutralization of the H4 K12 and K16 residues since the mobility of the acetylated and unmodified nucleosomes were the same (**Fig. 4d, lanes 1 and 2)**. Given that the mobility of a complex in a native PAGE gel depends on both the charge and the shape of the complex, the simplest interpretation of our results is that the BRD4-S/acetylated nucleosome complex is more compact than the BRD4-S/unmodified nucleosome complex.

Our structural and TR-FRET studies have pointed to three regions within or close to BRD4 BD1 that interact with the nucleosome: the BRD4 BD1 peptide binding pocket which interacts with acetylated H4 tail, the R68, K72 and K76 basic residues in BRD4 BD1 helix ⍺Z that interact with nucleosomal DNA, and the BRD4 basic patch 1 residues that appear to interact with the nucleosomal DNA minor groove near the nucleosomal dyad. We find that mutations in each of these three regions decreases the mobility of the BRD4-S/H4 K12acK16ac nucleosome complex on native PAGE. Both the Y97F,N140A mutations (m2 mutation) in the BRD4 BD1 peptide binding pocket and the R68E,K72E,K76E mutations in BRD4 BD1 ⍺Z (m28 mutation) result in BRD4/H4 K12acK16ac nucleosome complexes that migrate slower than the wild type complex (**Fig. 4d, lanes 6-8)**.

The effects of mutating BRD4 basic patch 1 residues were even more dramatic with retardation of the complex when the five basic patch 1 residues were mutated to Ala (m11 mutation) and even larger retardation when the five basic patch 1 residues were mutated to Glu (m12 mutation) (**Fig. 4d, lanes 8-10)**. This was true whether H4 K12acK16ac or unmodified nucleosomes were used (**Fig. 4d, lanes 11-13)**. The effects were specific since mutating the BRD4 BD1 K99 and K102 residues on the BD1 surface facing away from the nucleosome to Glu (m27 mutation) did not affect the mobility of the BRD4-S/nucleosome complex (**Fig. 4d, lanes 4 and 5)**. It is worth noting that the slower mobility of the Ala or Glu mutations is not simply the effect of changing the charge on BRD4-S. Since the Ala or Glu mutations make the BRD4-S protein more negatively charged, one might expect that the BRD4-S/nucleosome complex would migrate faster in a native DNA PAGE gel. However, the Ala or Glu mutations in BRD4 helix ⍺Z or basic patch 1 resulted in slower mobility BRD4-S/nucleosome complexes. To confirm that mobility shift is not an artifact of native gel electrophoresis, we have analyzed the elution of BRD4/H4 K12acK16ac nucleosomes by Superdex 200 Increase size exclusion chromatography (**Fig. 5**). We find that the BRD4-S/nucleosome complex containing the BRD4 basic patch 1 to Glu mutations elutes earlier than the wild type BRD4-S/nucleosome complex, indicative of a greater hydrodynamic radius. The BRD4 protein on its own elutes at essentially the same time with or without the basic patch 1 to Glu mutations. These results indicate that BRD4-S adopts a more compact structure on H4 K12acK16ac nucleosomes than on unmodified nucleosomes and that disrupting BRD4’s contacts with the nucleosome decreases the compactness of the BRD4/nucleosome complex.

**Figure 5:**
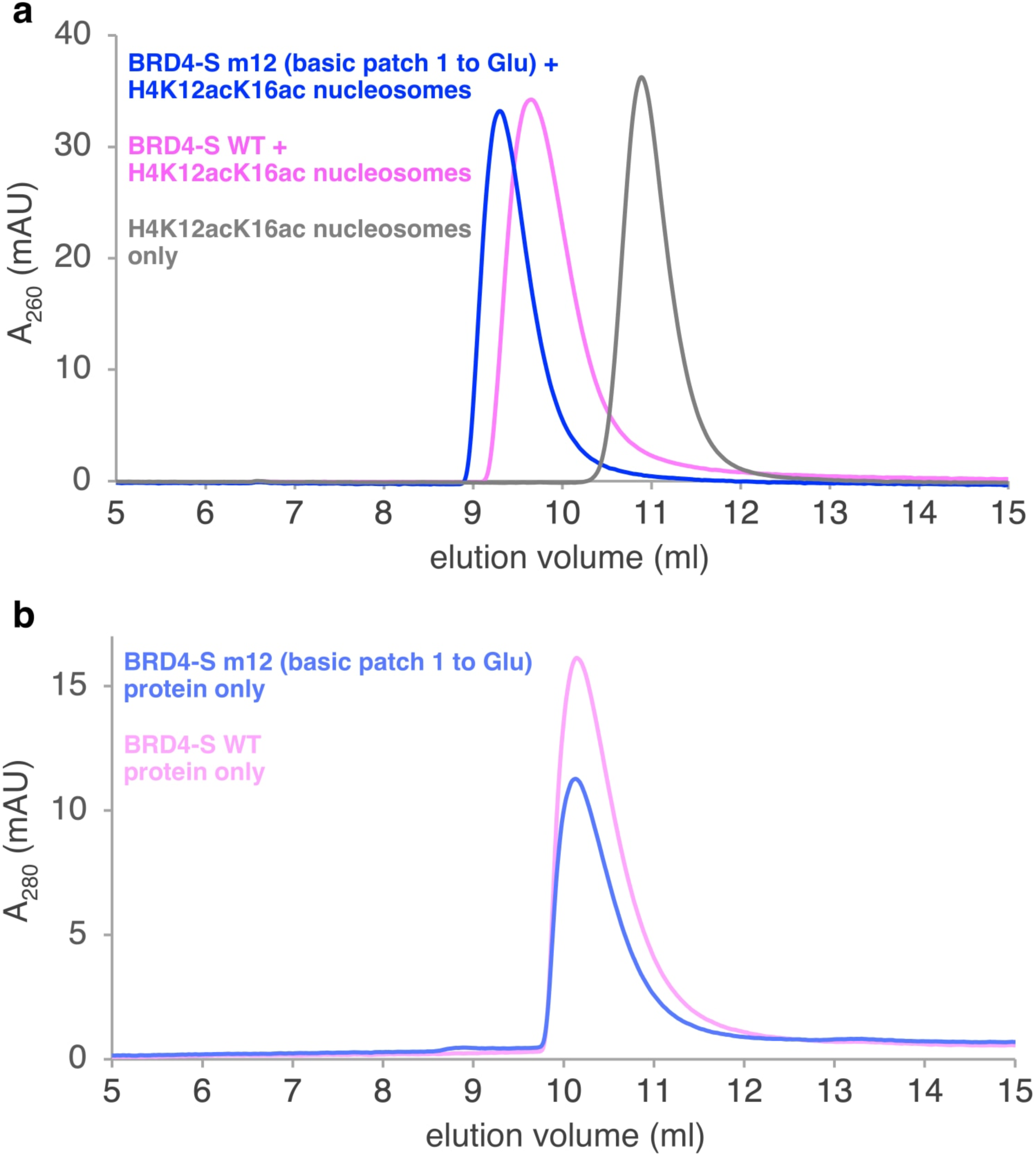
Superdex 200 increase size exclusion chromatography of BRD4-S/nucleosome complexes and BRD4-S proteins. (a) the BRD4-S m12/H4 K12acK16ac nucleosome complex (blue) elutes earlier than the BRD4-S WT/H4 K12acK16ac nucleosome complex, (b) the BRD4-S m12 protein elutes at similar time as the BRD4 WT protein.

BRD4 mutations in basic patches 2, 3 and 4 also decreased the mobility of the BRD4/nucleosome complex in native gel electrophoresis, although generally to a lesser degree than for basic patch 1 (**Fig. 6a, lanes 3-8)**. Similar effects were observed with H4 K12acK16ac and unmodified nucleosomes, consistent with binding to the acetylated H4 tail not being the major determinant in nucleosome binding by BRD4 (**Fig. 6a vs 6b)**. In contrast to the basic patches 1-4, mutating BRD4 basic patch 5 did not affect the mobility of the BRD4-S/nucleosome complex despite our finding that the same mutations severely affected nucleosome binding in the TR-FRET assay (**Fig. 6a, lane 10)**. This suggests that the compactness of the BRD4-S/nucleosome is not necessarily directly related to how tightly BRD4 binds to the nucleosome. The effect of mutating combinations of BRD4 basic patches appears to be additive in the gel mobility assay, with mutating basic patches 1, 2 and 3 or patches 4 and 5 having a larger retardation effect than individual patches (**Fig. 6a, lanes 11-12)**.

**Figure 6:**
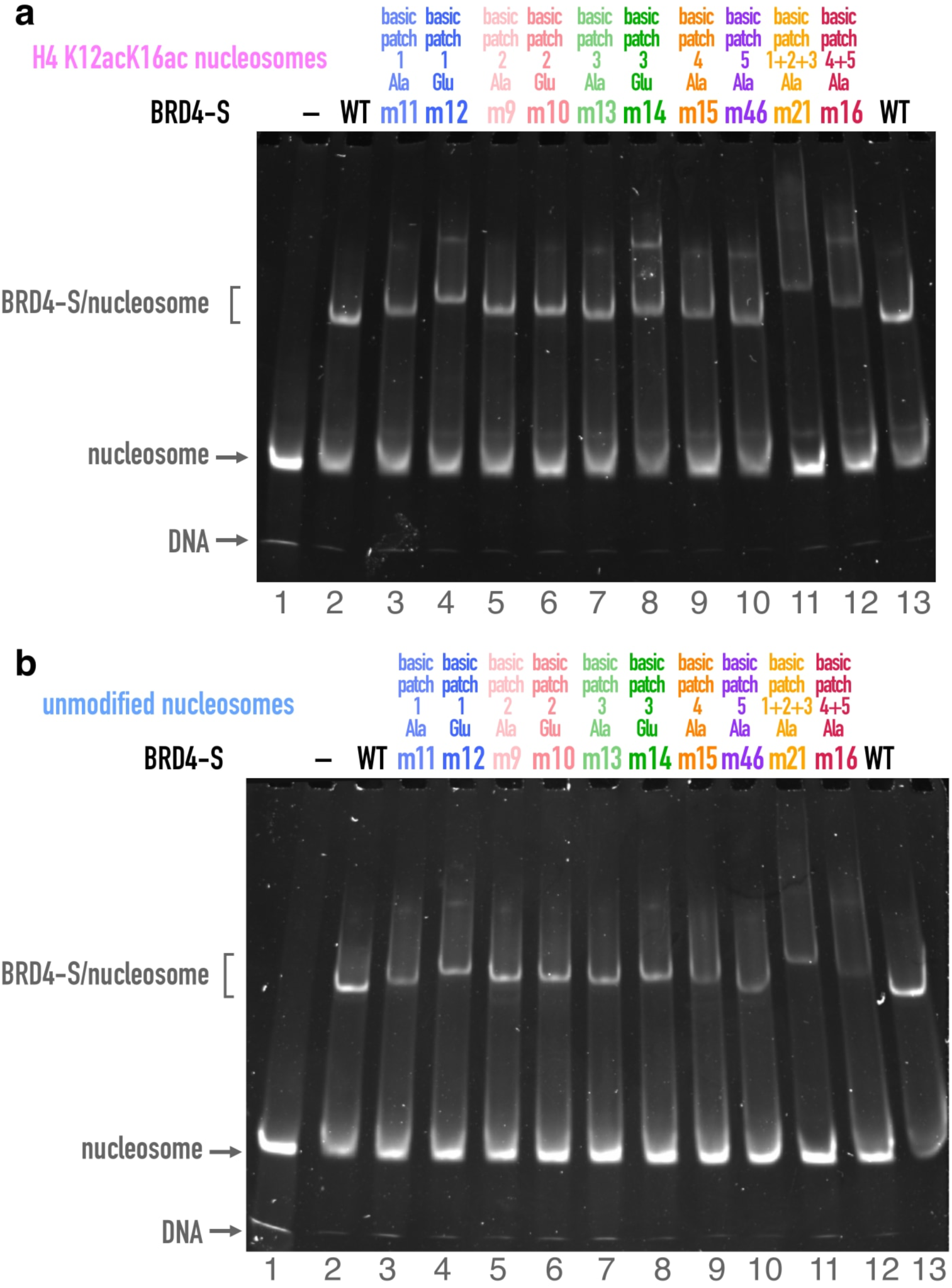
Effect of BRD4 basic patch mutations on the compactness of BRD4-S/nucleosome complexes in gel mobility shift assay. using (a) H4 K12acK16ac nucleosomes and (b) unmodified nucleosomes.

## Discussion

We show that BRD4 interacts with its nucleosome target via multivalent interactions involving multiple regions of BRD4 and the nucleosome besides the well-characterized BRD4 bromodomain binding to acetylated histone peptides. Our structural studies reveal that BRD4 BD1 is positioned on the nucleosome face to bind to nucleosomal DNA in addition to binding the acetylated H4 tail through the bromodomain peptide binding pocket. This nucleosomal DNA binding is consistent with previous studies of the related BRDT protein which highlighted a positively charged patch on its BD1 and nonspecific binding of that bromodomain to DNA^54^. BRD2, BRD3, BRD4 and BRDT all share a positively charged patch similarly localized on their BD1^54^ and they all bind to acetylated H4 tails using BD1^30,46,55^. Since the BRD4 residues R68, K72 and K76 that we observe to bind nucleosomal DNA appear to be part of this conserved positively charged bromodomain patch **(Extended Data Fig. 8**), it seems reasonable to anticipate that BD1 of BRD2, BRD3 and BRDT may bind in a similar orientation on the nucleosome as BRD4 does in our cryo-EM reconstruction. In contrast, since only the BRD2, BRD3, BRD4 and BRDT residues in BD2 corresponding to BRD4 K72 are positively charged **(Extended Data Fig. 8**), BD2 of the other BET proteins might not bind to the nucleosome in this orientation.

Although most inhibitors of BRD4 target the peptide binding pocket, a new class of BRD4 inhibitors has been found to bind to a distinctly different site. The ZL0590 BRD4 inhibitor which possesses anti-inflammatory activities has been determined to bind to BRD4 BD1 helices ⍺A and ⍺B^56^. Superposition of the BRD4 BD1/ZL0590 crystal structure with our BRD4/nucleosome cryo-EM structure shows that the ZL0590 compound is positioned away from the nucleosome (**Fig. 7**). If ZL0590 acts in the context of chromatin and ZL0590 does not act as an allosteric effector of acetyl-lysine peptide binding, the BRD4 BD1 ⍺A and ⍺B surface it binds to may be targets of effector proteins associated with its anti-inflammatory properties.

**Figure 7:**
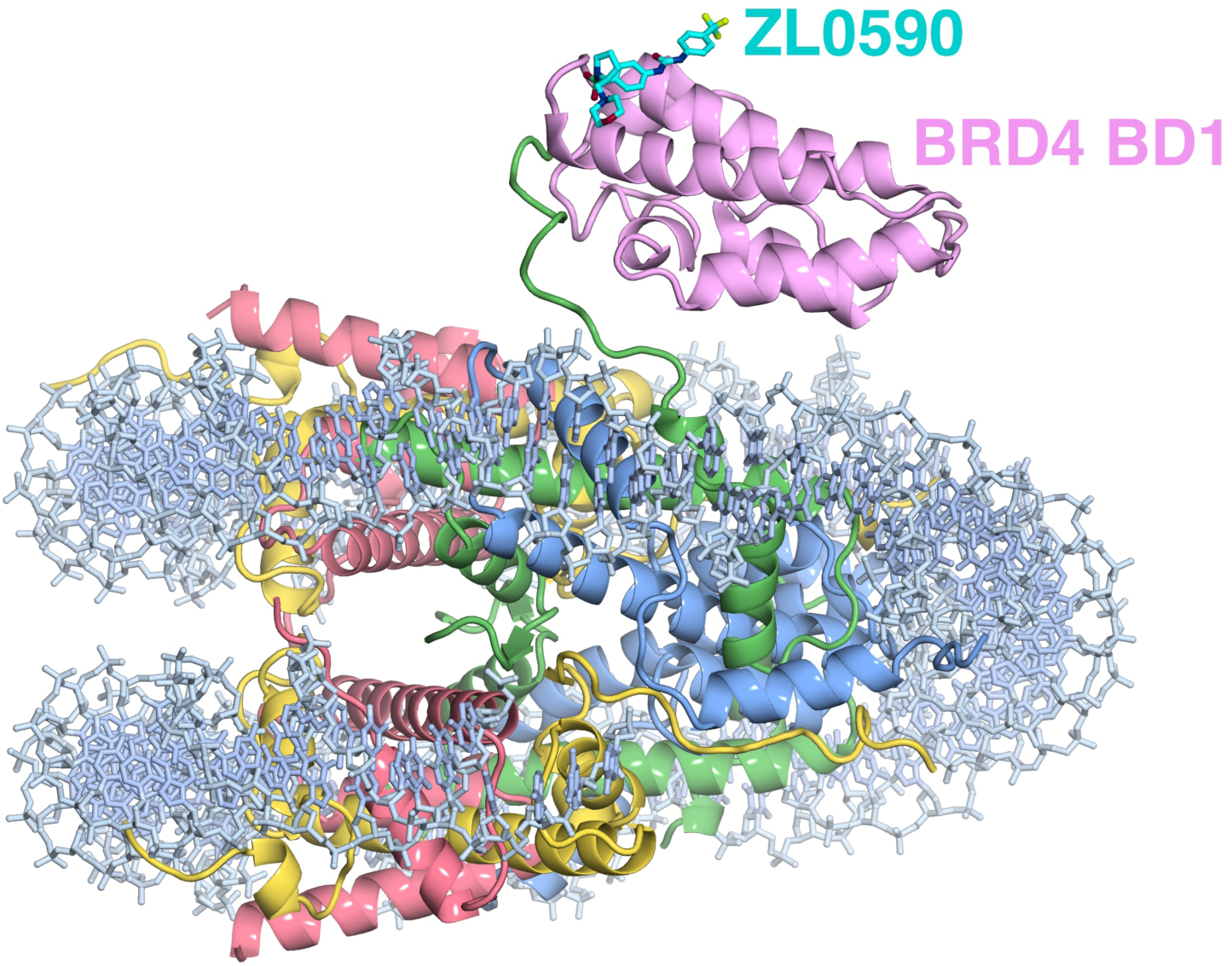
Structural model for ZL0590 inhibitor bound to the BRD4/nucleosome complex. Superposition of ZL0590/BRD4 BD1 crystal structure (PDB 6U0D) with BRD4/nucleosome structure (this work) via BRD4 BD1.

Based on the prevailing model that BRD4 is recruited to chromatin via bromodomain binding to acetylated histone tails, we had anticipated that BRD4 would require histone acetylation to bind to nucleosomes. Contrary to this expectation, we find that BRD4 binds tightly (10-40 nM dissociation constant) to both diacetylated and unmodified nucleosomes with only a 3 to 4-fold preference for diacetylated nucleosomes at 150 mM NaCl and even less preference at lower salt concentrations. Our mutational analyses indicate that critical interactions with the nucleosome are made by other BRD4 regions. We have identified five basic patches in BRD4 that appear to mediate redundant interactions with nucleosomal DNA. These basic patches likely bind nonspecifically to nucleosomal DNA in a conformationally heterogenous manner. This is consistent with the finding that residues in the BID region (which includes basic patches 4 and 5) help BRD4 bind DNA and native nucleosomes, and with NMR studies that a BRD4 BD1-BD2 construct binds DNA in a conformationally dynamic fashion^47,48^. The lack of a unique binding mode likely accounts for why the BRD4 basic region interactions with DNA were not observed in our cryo-EM maps.

We provide evidence that although BRD4 binds with similar affinity to diacetylated and unmodified nucleosomes, the resulting complexes are conformationally different. Experiments using the gel mobility shift assay and size-exclusion chromatography suggest that BRD4 adopts a more compact conformation on acetylated vs non-acetylated nucleosomes. Mutating the BD1 peptide binding pocket and the BD1 nucleosome DNA-binding surface had similar effects in the gel mobility shift assay as removing the histone H4 acetylation marks, consistent with these BRD4 BD1 regions interacting with the nucleosome. Mutating BRD4 basic patches 1-4 also appear to make the corresponding BRD4/nucleosome complexes less compact, with mutations in basic patch 1 immediately following BD1 having the largest effect.

A mechanism of the role of basic patch 1 in nucleosome binding is suggested by our cryo-EM maps which show that the BRD4 region immediately C-terminal to BD1 approaches the nucleosomal DNA near the dyad. This positions the basic patch 1 to enter the DNA minor groove. We propose that the two overlapping RGR sequences enable redundant AT-hook-like binding in the DNA minor groove. Doing so provides an additional tether of BRD4 BD1 to the nucleosome in addition to the bromodomain-acetylated H4 tail and the bromodomain ⍺Z-nucleosome DNA interactions. Disrupting any of these tethers destabilizes the positioning of BRD4 BD1 on the nucleosome, resulting in a conformationally less compact and presumably more flexible complex. The finding that mutations of the basic residues in the basic region 1 have a stronger effect on the affinity of nucleosome binding and on the extent of the gel mobility shift underlines the importance of basic region 1’s interactions with the nucleosome, perhaps to help position BD1 appropriately on the nucleosome face. The interactions of BD1 with the nucleosome do not involve a large contact surface area to stabilize the binding (occluded solvent accessible surface area of 1450 Å^2^ vs 2725 Å^2^ for SIRT6/nucleosome or 3570 Å^2^ for SIR3/nucleosome^39,57^ for example). Instead, the three contact points (BD1 binding pocket with acetylated histone H4 tail, BD1 ⍺Z interactions with nucleosomal DNA and basic patch 1 interactions with nucleosomal DNA at the dyad) tether BD1 to the nucleosome without a large contact surface. The potential resulting wobbliness of BD1 on the nucleosome may account for the lower resolution of cryo-EM reconstruction for the bromodomain component of the complex compared to the nucleosome.

The interactions of the BRD4 basic patch 1 with the nucleosome are particularly interesting because they suggest a mechanism for regulating the recruitment of BRD4 to chromatin. BRD4 has been found to be methylated at R179, R181 and R183, the very arginine residues in the RGRGR overlapping RGR sequence in basic region 1 that we have identified as potentially interacting with the DNA minor groove at the nucleosome dyad^58–60^. In one study, methylation of these three BRD4 arginine residues by the arginine methyltransferases PRMT2/4 regulated transcription and DNA repair^58^. In a second study, methylation of the same three BRD4 arginine residues promotes BRD4 phosphorylation and ovarian cancer metastasis^59^. More recently, methylation of these three BRD4 arginine residues by PRMT1 was found to promote partial epithelial-mesenchymal transformation and renal fibrosis^60^. Our findings which suggest that Arg179, Arg181 and Arg183 stabilize BRD4 interaction to the nucleosome by binding in the DNA minor groove could provide the molecular basis for how arginine methylation affects BRD4 function through its binding to the nucleosome. Further studies will be needed to examine how symmetric arginine methylation by PRMT2/4 and asymmetric arginine methylation by PRMT1 affect BRD4 binding to the nucleosome.

Our unexpected finding that BRD4 binds tightly to unmodified nucleosomes and that H4 K12 and K16 acetylation improves binding affinity only 3-4 fold at 150 mM NaCl concentration shows that BRD4 does not require histone acetylation to bind to nucleosomes. This is contrary to the expectation that histone acetylation is required for BRD4 to bind to chromatin based on observations that BRD4 binds specifically to acetylated chromatin in cells^26^. It is, however, consistent with findings that the JQ1 compound, which competes for acetylated histone tail binding in the bromodomain peptide binding site, only partially removed BRD4 from chromatin *in vivo*^21^. Nevertheless, our finding that BRD4-S binds tightly to nucleosomes with ∼40 nM dissociation constant in the absence of acetylation begs the question of how BRD4 distinguishes between unmodified and acetylated chromatin in the cell. There are at least three possible resolutions to this apparent paradox. Firstly, BRD4 could bind differently to nucleosome arrays than it does to the mononucleosome substrates used in this study. Secondly, BRD4 in a cell might contain post-translational modifications, including phosphorylation, which might modulate its binding to chromatin^45^. Thirdly, BRD4 has been shown to form biomolecular condensates providing different environments from our in-solution experiments^47,61,62^. In any case, our results that BRD4 binds tightly to unmodified nucleosomes does suggest that specific recruitment of BRD4 to acetylated nucleosome in cells may involve mechanisms that prevent BRD4 from binding to unmodified nucleosome as well as ones that promote binding to acetylated nucleosomes. Our results also provide more information to understand how targeted BRD4 degradation can inhibit cancer more effectively than BET inhibition^19,63^.

In summary, our structural and biochemical studies of how BRD4 interacts with acetylated nucleosomes challenge the canonical view that BRD4 is recruited to chromatin via bromodomain-acetylated histone interactions. We find instead that BRD4 utilizes multiple surfaces of BD1 as well as basic regions 1-5 to bind to H4 K12acK16ac nucleosomes. The interactions with nucleosomal DNA observed structurally for BD1 and surmised for the basic regions 1-5 appear to be responsible for BRD4’s ability to bind to nucleosomes with high affinity even in the absence of histone modifications. In contrast, the BRD4 bromodomain-acetylated histone interaction contributes only a modest 3- to 4-fold to the binding affinity to nucleosomes. Disrupting the BD1-acetylated histone H4 interactions and the presumed basic patch-DNA interactions does, however, adversely affect the conformational compactness of the BRD4/nucleosome complex. The functional significance of the conformationally compact BRD4/acetylated nucleosome complex remains to be determined. It is possible that the compact BRD4/acetylated nucleosome conformation provides a combination of surfaces that is recognized by a BRD4-interacting cofactor, thus ensuring binding of this cofactor specifically to a BRD4/acetylated nucleosome and not to BRD4/unacetylated nucleosome. Additional *in vitro* and *in vivo* studies will be necessary to test this hypothesis.

## Materials and Methods

### Protein purification

The coding region of the short isoform of human BRD4 (residues 1–722) was amplified from the plasmid pGEX-6P-1 BRD4 full-length (a gift from Peter Howley, AddGene plasmid #14447^64^) and cloned into the pST50Tr-HISN expression vector^65^. Point mutants were generated using PCR-based site-directed mutagenesis. HIS-tagged BRD4 was expressed in *E. coli* BL21(DE3)pLysS cells through IPTG induction or autoinduction^66^ at 23°C and purified from crude lysates via metal affinity chromatography (Talon resin, Clontech). For structural studies, the HIS tag was removed using tobacco etch virus (TEV) protease, while it was retained for TR-FRET studies. Further purification was achieved using Source S cation-exchange chromatography (Cytiva) followed by Source Q anion-exchange chromatography (Cytiva). Dynamic light scattering analysis of BRD4 proteins confirmed the absence of aggregation.

### Nucleosome preparation

Recombinant *Xenopus laevis* and human core histones, as well as nucleosome core particles, were prepared following previously described protocols^67^. Nucleosomes contained *Xenopus* histones H3 (98.5% identical to human H3) and H4 (identical to human H4) as well as human H2A and H2B. All nucleosomes were purified using SourceQ anion-exchange liquid chromatography. Acetylated histone H3 and H4 variants were expressed in *E. coli* through amber suppression acetyl-lysine incorporation and purified via metal affinity chromatography (Ni-NTA Superflow resin, QIAGEN)^68,69^. Diacetylated histone H4 was treated with TEV protease to remove protein tags^69^. The tailless histone H4 protein corresponds to H4(24-102). Histone mutations were generated using PCR-based site-directed mutagenesis. The H4 K12acK16ac modifications were validated by mass spectrometry and Western blotting **(Extended Data Fig. 9**). Nucleosomes for cryo-EM studies contained a 165 bp (10+145+10) Widom 601 nucleosome positioning sequence.

### Cryo-EM sample preparation and data collection

BRD4-S was reconstituted with nucleosomes in reconstitution buffer [10 mM HEPES pH 7.5, 20 mM KCl, and 1 mM dithiothreitol (DTT)] at a BRD4-S to nucleosome ratio of 2.1:1. The resulting complex was cross-linked using the GraFix method ^70^. A glycerol gradient was prepared using a light buffer [10 mM HEPES pH 7.5, 20 mM KCl, 1 mM DTT, and 10% glycerol] and a heavy buffer [10 mM HEPES pH 7.5, 20 mM KCl, 1 mM DTT, 40% glycerol, and 0.15% glutaraldehyde], generating a 10–40% glycerol gradient with a corresponding 0–0.15% glutaraldehyde gradient. Gradient fractions were analyzed via native polyacrylamide gel electrophoresis (PAGE), and fractions containing the BRD4/nucleosome complex were concentrated to ∼4 μM.

Cryo-EM grids of the BRD4/nucleosome complexes were prepared using standard procedures^71^. A 3 μl aliquot of the BRD4/nucleosome sample was applied to holey carbon 1.2/1.3 Cu300 mesh grids (Quantifoil) using a FEI Vitrobot Mark IV, maintained at 4°C with 100% humidity. The sample was blotted for 3.5 seconds with a blot force of −1 and subsequently plunge-frozen into liquid ethane.

The microscopy dataset was collected at the Pacific Northwest Cryo-EM Center using a FEI Titan Krios microscope operated at 300 keV and equipped with a Gatan K3 direct electron detector. Two datasets, consisting of 11,012 and 18,227 movies, respectively, were combined for a total of 29,239 movies. Data were collected in counting mode at a magnification of ×22,500, corresponding to a pixel size of 1.0125 Å. Images were recorded with a defocus range of −0.5 to −2.0 μm and an accumulated electron exposure of ∼40 e⁻/Å² distributed across 40 frames **(Extended Data Table 1**).

### Cryo-EM data processing

The BRD4/nucleosome dataset was processed in cryoSPARC ^72^. Raw movies were motion-corrected using Patch Motion Correction (multi), and dose-weighted micrographs^73^ were generated. Defocus values were estimated with Patch CTF Estimation (multi). Initial particle picking was performed on a subset of micrographs using Blob Picker, with the particle diameter set to 100–200 Å. Picked particles were extracted with 4× binning in a 256-pixel box Fourier-binned to 64 pixels, resulting in a pixel size of 4.236 Å. Two-dimensional (2D) classification was conducted to remove classes containing obvious junk (Extended Data Fig. 2).

Remaining particles were subjected to *ab initio* reconstruction to generate input classes for heterogeneous refinement. A cleaned subset of nucleosomal particles was then used to generate templates for particle picking on the complete dataset. Template Picker, with a particle diameter set to 200 Å, was employed to obtain particles across all micrographs. Subsequent rounds of 2D classification removed additional junk classes, and *ab initio* reconstruction was repeated to refine input classes for heterogeneous refinement.

Multiple rounds of 3D classification on a cleaned dataset of nucleosomal particles ultimately yielded a single class with clear density for the BRD4 bromodomain. To improve the quality of the complex, focused 3D classification was performed with a mask centered on the BRD4 bromodomain. From this process, a class of 80,704 particles, representing the most stable positioning of BRD4 bromodomain I on the nucleosome, was selected. These particles were further refined using non-uniform refinement in cryoSPARC^74^.

### Model building and refinement

The nucleosome component of the complex was modeled using Protein Data Bank (PDB) entry 3LZ0^75^ as the starting model, while PDB entry 3UVX ^46^ served as the starting model for BRD4 BD1. Both models were rigid-body fitted into the 2.89-Å reconstruction using the “fit in map” function in UCSF ChimeraX^76^ and subsequently optimized in Coot^77^.

The highest-resolution 2.89-Å reconstruction was used in Coot to refine the histones and core DNA, and to build the histone H4 N-terminal tail. The final model was refined with *phenix.real_space_refine*^78^, incorporating secondary structure, Ramachandran, and rotamer restraints. The model underwent manual validation in Coot and comprehensive validation (cryo-EM) in Phenix using MolProbity^79^. Model statistics are provided in **Extended Data Table 2**.

### Time-resolved FRET nucleosome binding assays

TR-FRET assays were performed as described previously^43^. Assay buffer (20 mM HEPES pH 7.5, 70-150 mM NaCl, 5 mM DTT, 5% glycerol, 0.01% NP-40, 0.01% CHAPS, 0.1 mg/mL BSA) was used to prepare 2x acceptor mixtures containing ULight anti-6xHIS acceptor antibody (Revvity) with 6xHIS-tagged BRD4 proteins at a ratio of 1:4 or 1:8 and serially diluting across 12 concentrations. 2x donor mixtures were prepared by mixing 4 nM streptavidin-Eu (Revvity) with or without 2 nM nucleosomes containing 177 bp of Widom 601 DNA (31+145+1) with a 5′ biotin group on the 31-bp extension. When used, the JQ1 compound was present at a concentration of 20 µM.

Samples were prepared by mixing 5 μL of donor mixtures with 5 μL of acceptor mixtures at each concentration in white, non-binding 384-well plates. Following a 30-minute incubation, fluorescence signals were acquired at room temperature in a Victor Nivo multimode fluorescent plate reader (PerkinElmer) using an excitation filter at 320 nm and emission filters at 615 and 665 nm. Emission signals were measured simultaneously following a 100-μs delay. Binding data were fit to the Hill equation:

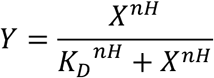

where Y is the normalized fluorescence signal, X is the acceptor protein concentration, K_d_ is the apparent dissociation constant, and nH is the Hill coefficient^80^. Data were fit using the “curve_fit” function of the SciPy Python library^81^. Data are plotted as means ± standard deviations of triplicate measurements, and K_d_ and Hill coefficient values are reported as means ± standard error of the mean from independent fits. We estimate we are able to detect nucleosome binding with dissociation constants less than 20 µM.

### Electrophoresis gel mobility shift assay

20 µl binding reactions were performed with 100 nM BRD4 variants incubated with 100 nM nucleosome core particles for 30 minutes at room temperature. The binding buffer consisted of 20 mM HEPES pH 7.5, 125 mM NaCl, 5 mM DTT, 5% glycerol, 0.01% NP-40, 0.01% CHAPS, and 100 μg/ml BSA. Following incubation, the reactions were resolved on 4.5% native PAGE gels at 120 V for 40 minutes at room temperature. Gels were stained with ethidium bromide and imaged using a GelDoc Go system (Bio-Rad).

### Size exclusion chromatography

BRD4 and its mutants were reconstituted with nucleosomes in a buffer containing 10 mM HEPES (pH 7.5), 100 mM NaCl, 1 mM DTT, and 0.1 mM PMSF before injecting onto a Superdex 200 Increase 10/300 GL column (Cytiva) equilibrated with the reconstitution buffer and eluted at 0.5 ml/min. A 300 µL sample of the BRD4-nucleosome complex or nucleosome alone at a concentration of ∼1 µM was loaded onto the column and eluted using the same buffer.

## Acknowledgements

We thank George Loukopoulos for technical assistance, the Huck Institutes Cryo-Electron Microscopy Facility for use of the Talos Arctica G2 TEM and Joseph Cho for assistance with screening and data collection, Tara Fox at NCI, and Marzia Miletto at PNCC for assistance with EM data collection. We also thank the Tan Laboratory and the Penn State Center for Eukaryotic Gene Regulation for helpful discussions. We are grateful to Heinz Neumann and Jason Chin for sharing plasmid reagents used to prepare acetylated histones.

This work was supported by National Institutes of Health (NIH) grant R35 GM127034 (S.T.), NIH grant T32 G125592 to E.M.L., and NIH grants 1R01CA251698-01 and 1R01CA288743-01A1 to C.-M.C.

Research reported in this publication was supported by the Office of the Director, NIH, under award number S10OD026822-0. This project is funded, in part, under a grant from the Pennsylvania Department of Health using Tobacco CURE Funds. The Department specifically disclaims responsibility for any analyses, interpretations or conclusion. This research was, in part, supported by the National Cancer Institute’s National Cryo-EM Facility at the Frederick National Laboratory for Cancer Research under contract HSSN261200800001E. A portion of this research was supported by NIH grant R24GM154185 and performed at the Pacific Northwest Center for Cryo-EM (PNCC).

Molecular graphics and analyses were performed with UCSF ChimeraX, developed by the Resource for Biocomputing, Visualization, and Informatics at the University of California, San Francisco, with support from National Institutes of Health R01-GM129325 and the Office of Cyber Infrastructure and Computational Biology, National Institute of Allergy and Infectious Diseases.

## Author Contributions

J.Z. and S.T. constructed expression plasmids. J.Z., E.M.L., E.N.O and B.P.M. purified recombinant proteins and nucleosomes. J.Z. prepared cryo-EM samples and performed electrophoresis gel mobility shift and SEC assays. J.Z. performed cryo-EM analysis and model building, assisted by E.M.L. and J-P.A.. E.M.L. performed TR-FRET assays. S.-Y. Wu, C.-M. C. and S.T. conceived the project. J.Z., E.M.L. and S.T. planned the project, designed the experiments and wrote the manuscript. All authors discussed the results and commented on the manuscript.

## Data availability

The atomic coordinates of the BRD4-S/acetylated nucleosome and BRD4-S with basic patch 1/acetylated nucleosome complexes have been deposited to the RCSB Protein Data Bank with PDB ID XXXX and PDB XXXY. The cryo-EM Coulomb potential maps were deposited in the Electron Microscopy Data Bank as EMD-XXXXX (BRD4-S/acetylated nucleosome) and EMD-XXXXY (BRD4-S with basic patch 1/acetylated nucleosome). The raw cryo-EM data was deposited in EMPIAR (EMPIAR-ZZZ).

---

**Extended Data Figure 1:**
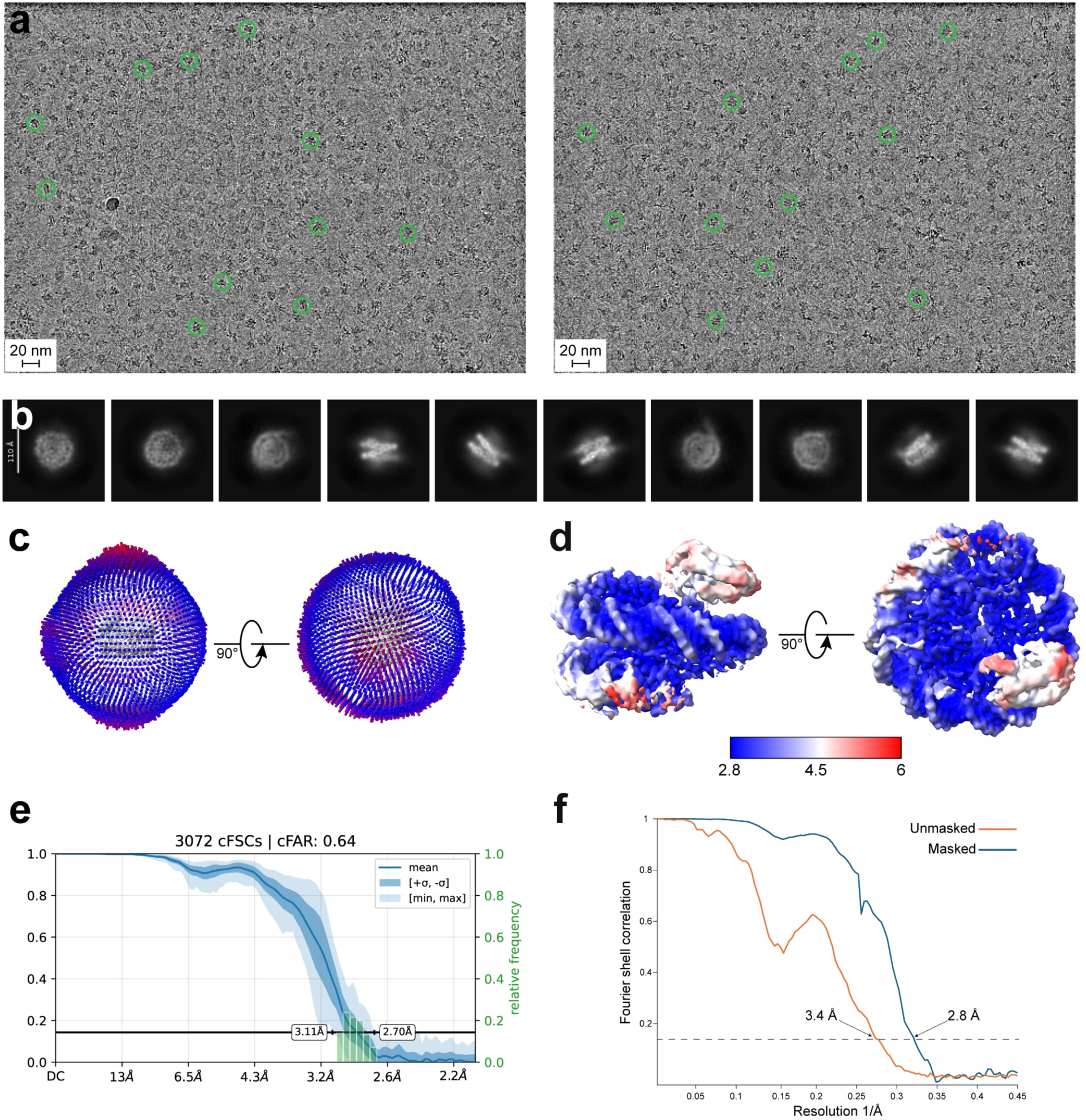
Cryo-EM studies of BRD4/nucleosome complex. (a) representative motion-corrected micrographs, (b) representative 2D classes, (c) angular distribution of particles used to generate the cryo-EM map, (d) cryo-EM map of the BRD4/nucleosome complex colored by estimated local resolution determined with FSC = 0.143 cutoff in cryoSPARC, (e) conical Fourier shell correlation (cFSC) curve of the BRD4-nucleosome structure at 2.89 Å resolution, calculated between two independent half-maps using a conical mask with a specified half-angle and axis in Fourier space in cryoSPARC. Lines and arrows indicate the axis of rotation between successive views, (f) unmasked (red) and masked (blue) Fourier shell correlation (FSC) curves between two independent half-maps determined in cryoSPARC.

**Extended Data Figure 2:**
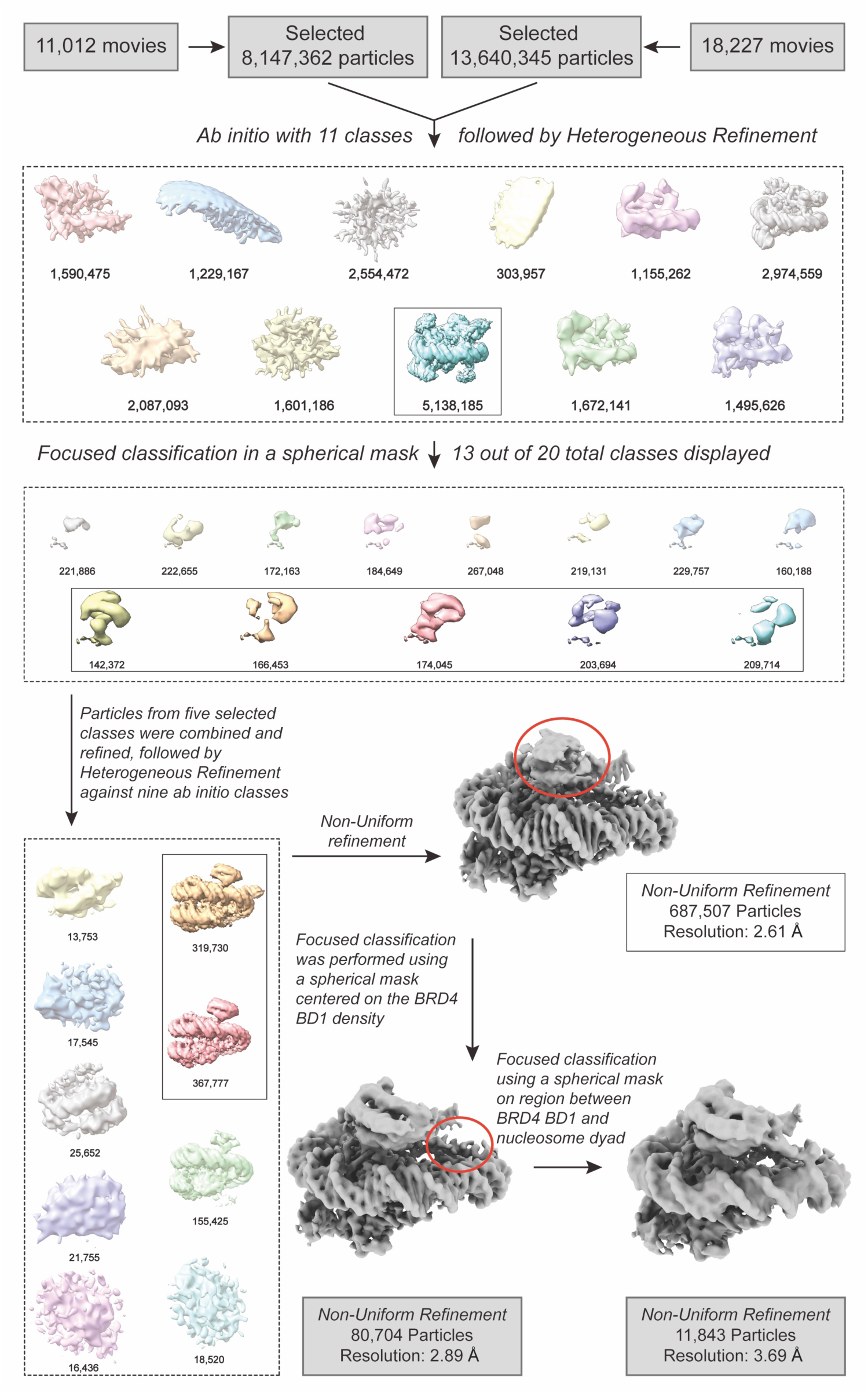
Cryo-EM data processing flow chart for BRD4-S/nucleosome complex. Schematic representation of the cryo-EM data processing pipeline for the BRD4/nucleosome complex, as described in the Methods section.

**Extended Data Figure 3:**
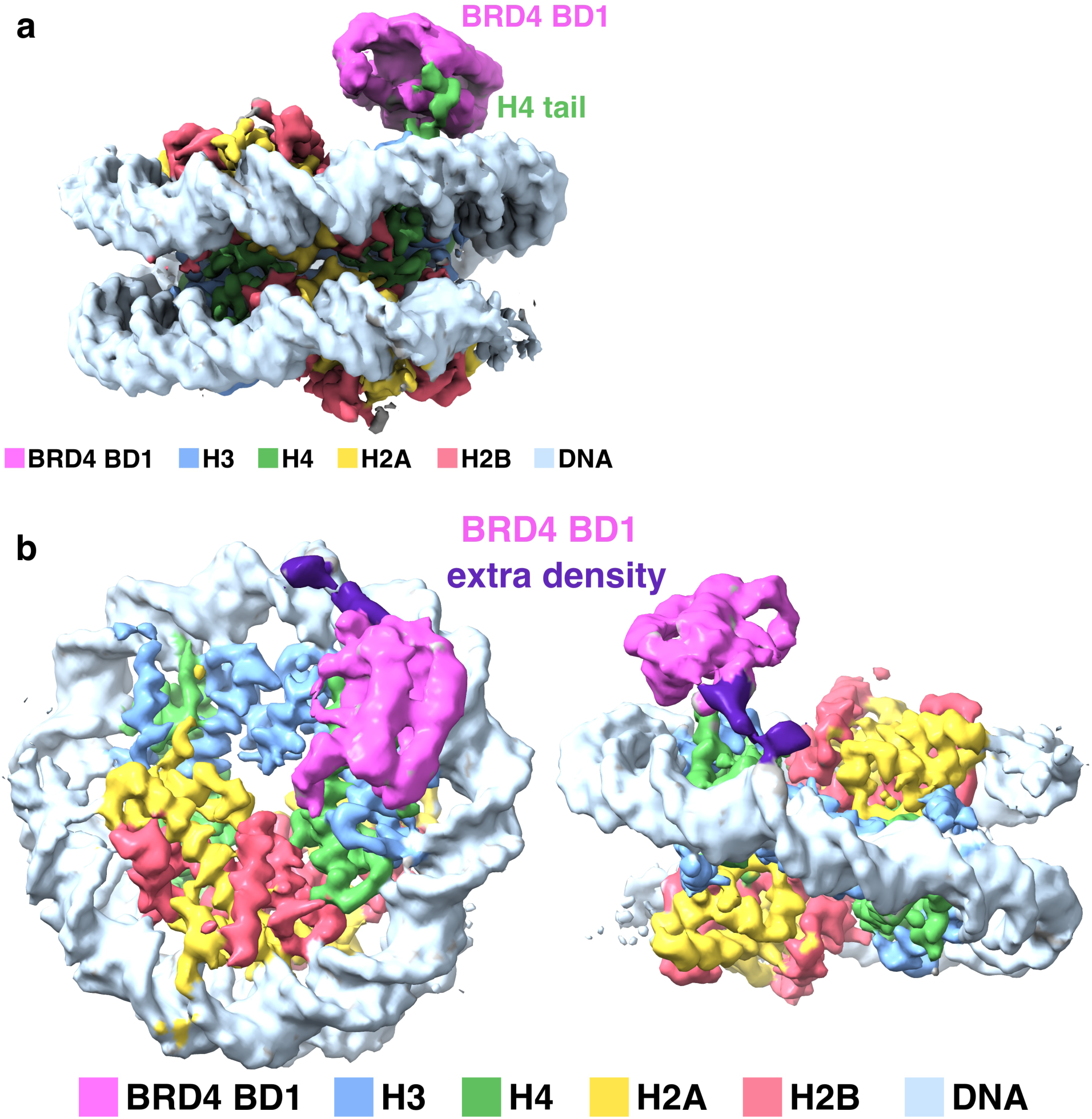
Cryo-EM maps of BRD4/nucleosome complex. (a) cryo-EM map of Fig. 1(a) viewed from the side to show interaction of histone H4 tail with BRD4 BD1, (b) cryo-EM map prepared from subset of particles shows extra density (purple) beyond the C-terminus of BRD4 BD1 (top view on left, side view on right).

**Extended Data Figure 4:**
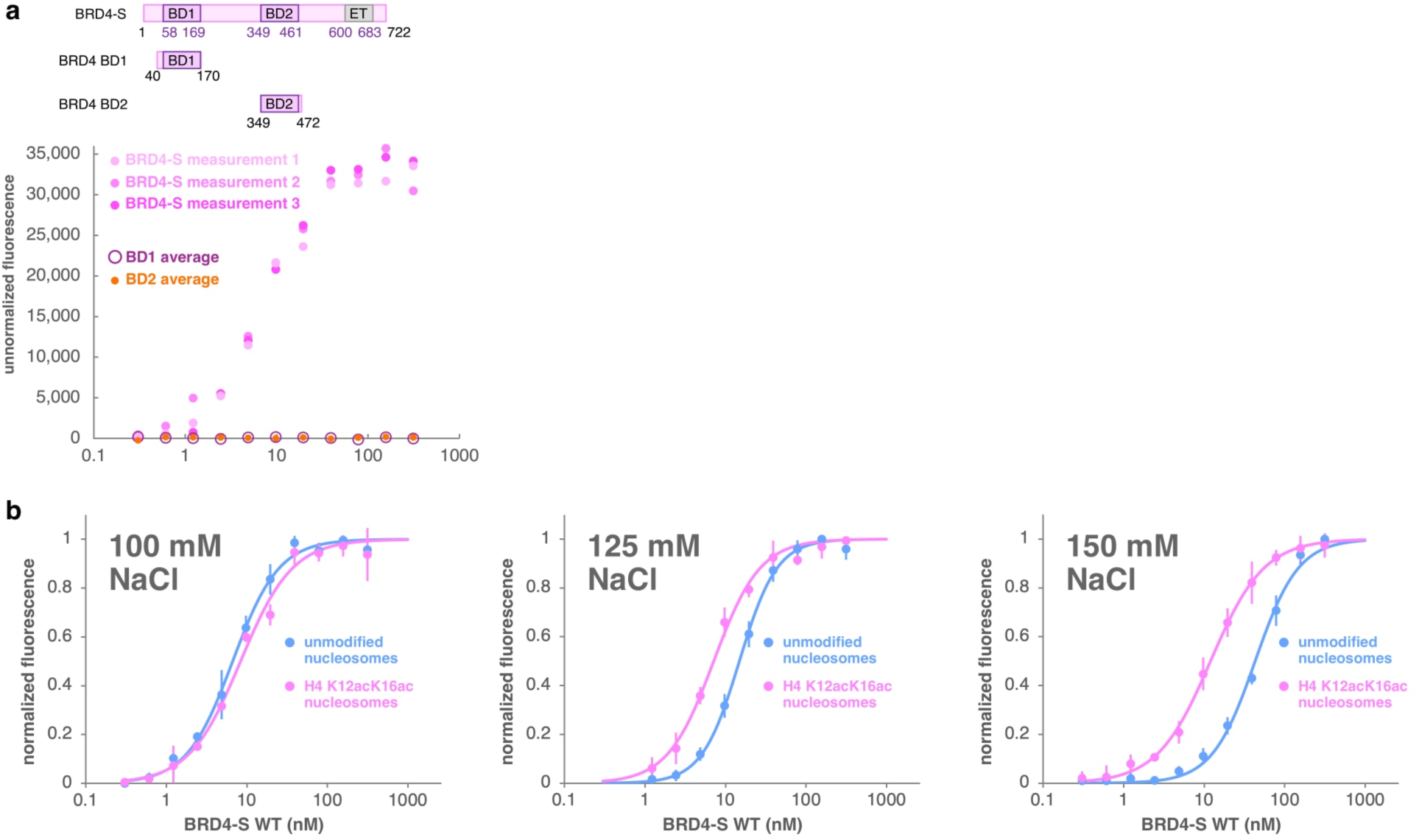
TR-FRET nucleosome binding assay. (a) top: cartoon representation of BRD4 constructs, bottom: individual TR-FRET unnormalized fluorescence (not normalized to maximum fluorescence) binding results of Fig. 2a for wild-type BRD4-S binding to H4 K12acK16ac nucleosomes and average of 3 measurements for BRD4 BD1 and BD2 binding to H4 K12acK16ac nucleosomes, (b) TR-FRET binding curves for BRD4-S binding to unmodified (blue) or H4 K12acK16ac (pink) nucleosomes in 100, 125 and 150 mM NaCl.

**Extended Data Figure 5:**
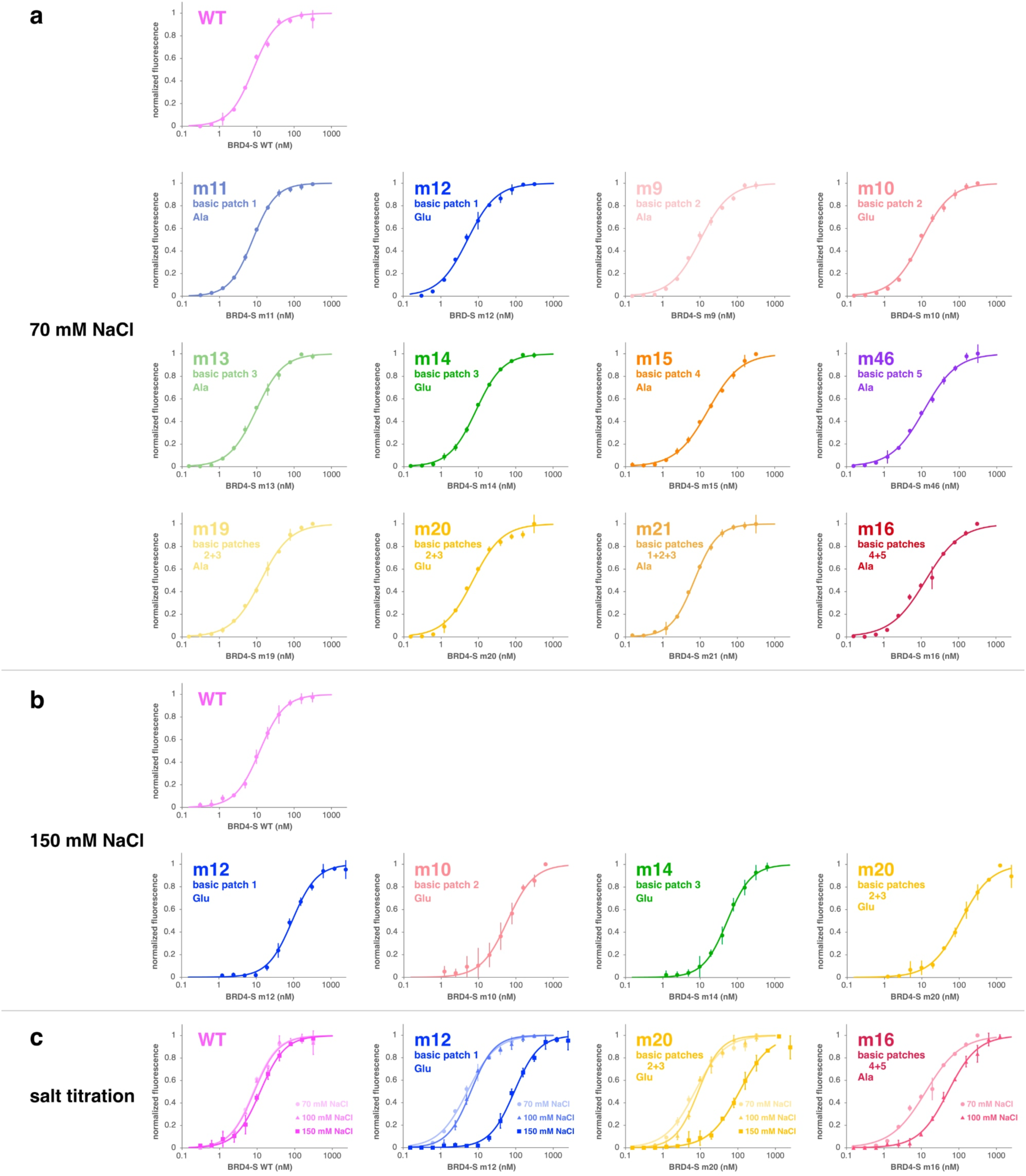
TR-FRET binding curves for BRD4-S basic patch mutants binding to H4 K12acK16ac nucleosome. (a) individual plots for binding in 70 mM NaCl (corresponds to Fig. 3b), (b) individual plots for binding in 150 mM NaCl (corresponds to Fig. 3c), (c) individual plots for salt titrations (corresponds to Fig. 3d)

**Extended Data Figure 6:**
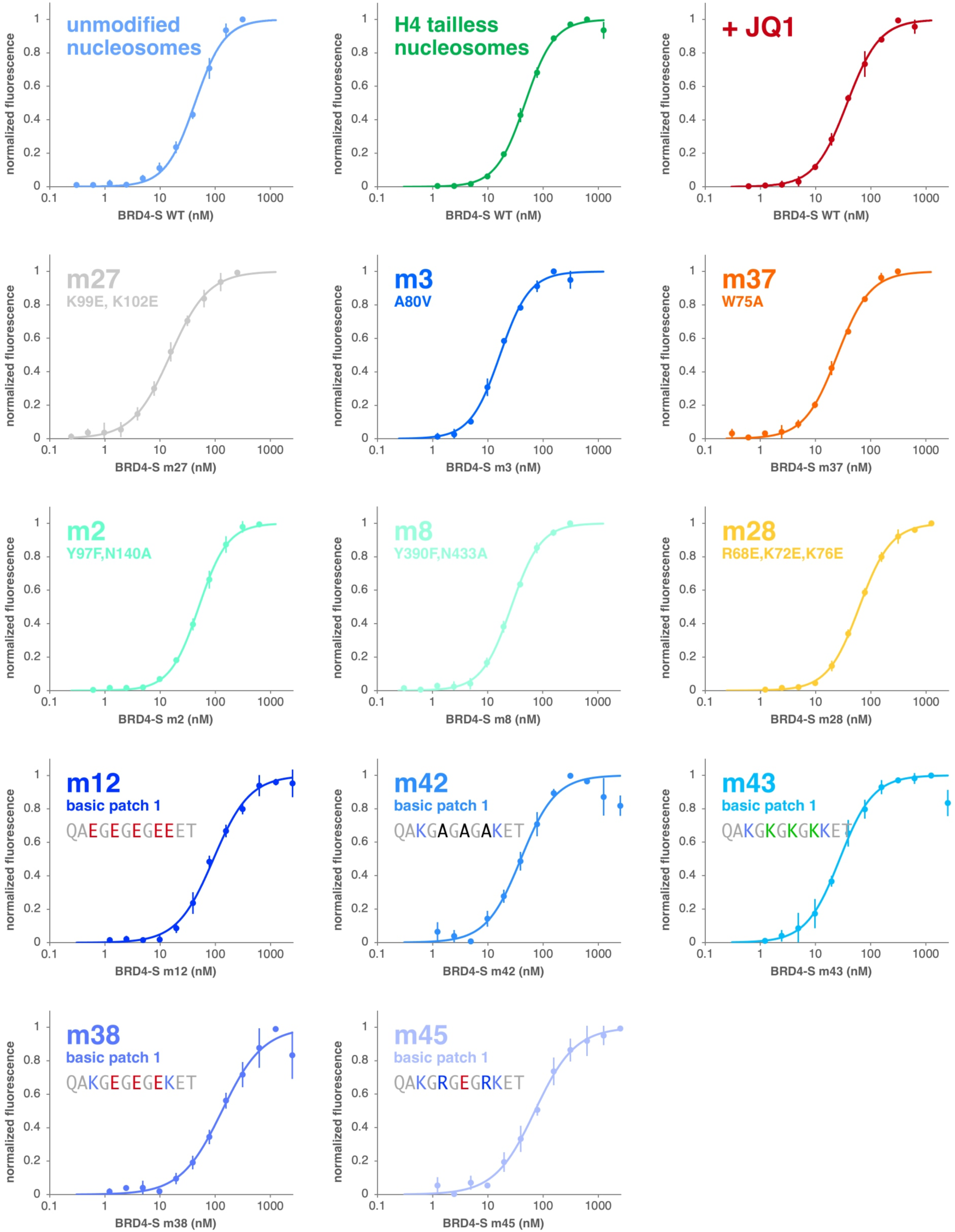
TR-FRET binding curves for BRD4-S BD1 mutants binding to nucleosomes. (corresponds to Fig. 4b and f). H4 K12acK16ac nucleosomes were used unless noted otherwise.

**Extended Data Figure 7:**
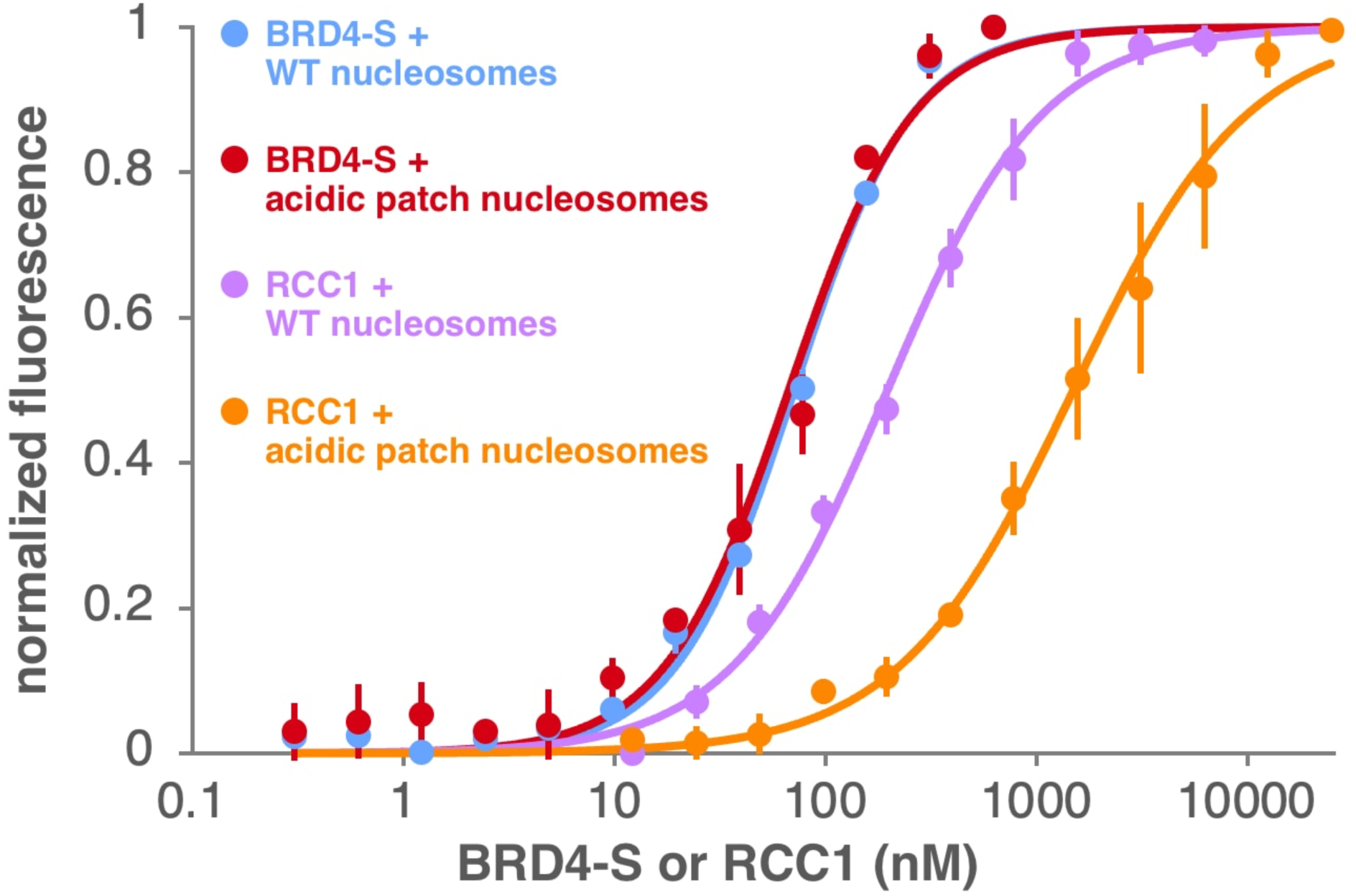
TR-FRET binding studies show BRD4-S binding to unmodified nucleosomes is unaffected by mutations to the nucleosome acidic patch. Acidic patch nucleosomes contain the H2A(E61A,E64A,D90A,E92A) quadruple mutation. BRD4-S binds wild-type nucleosomes (blue) and acidic patch nucleosomes (red) with similar affinity, but the RCC1 chromatin factor shown to use an arginine anchor to bind to the nucleosome acidic patch is adversely affected by the nucleosome acidic patch mutations.

**Extended Data Figure 8:**
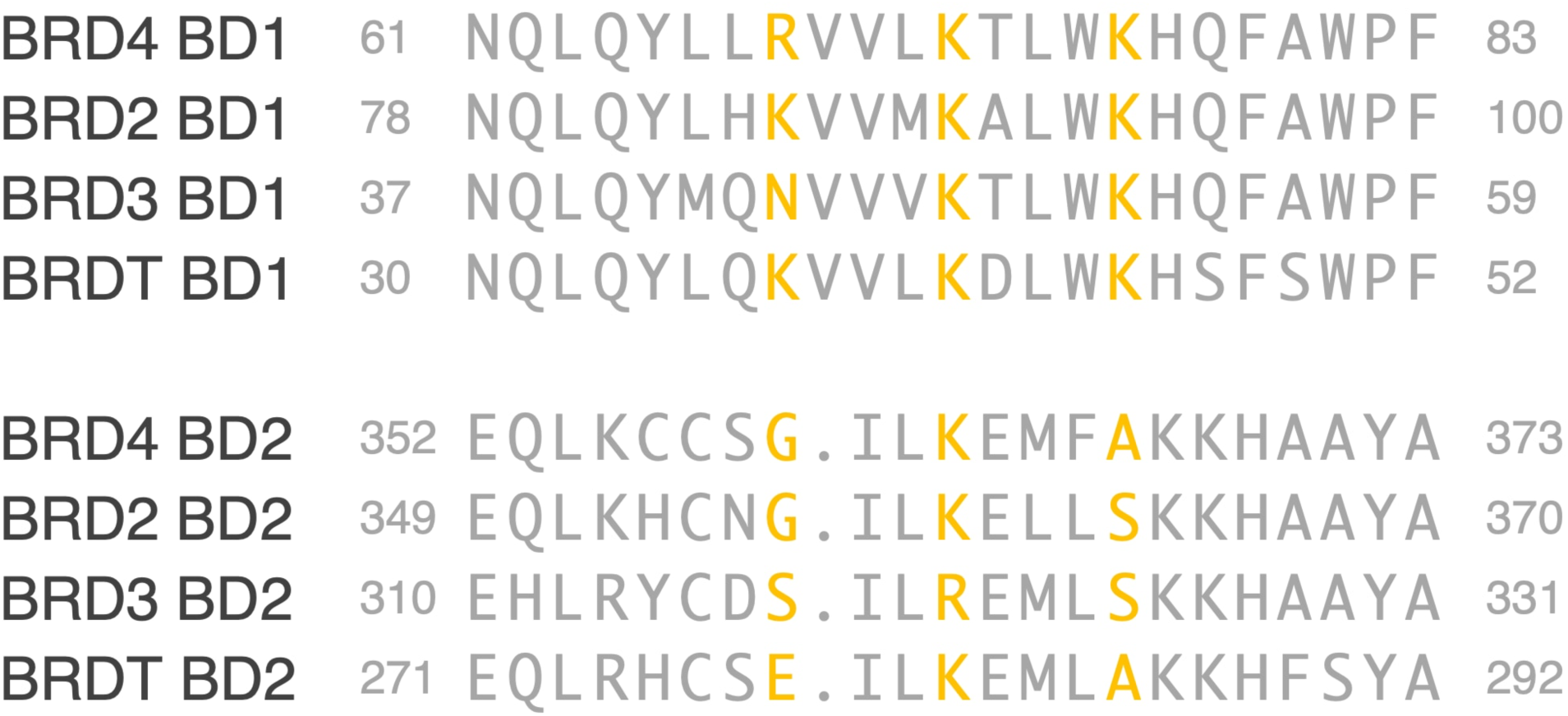
Sequence alignments of human BRD bromodomain 1 and 2 around the ⍺Z helix. The BRD4 residues R68, K72, K76 observed to interact with nucleosomal DNA and the BRD2, BRD3 and BRDT equivalent BD1 and BD2 residues are highlighted in yellow.

**Extended Data Figure 9:**
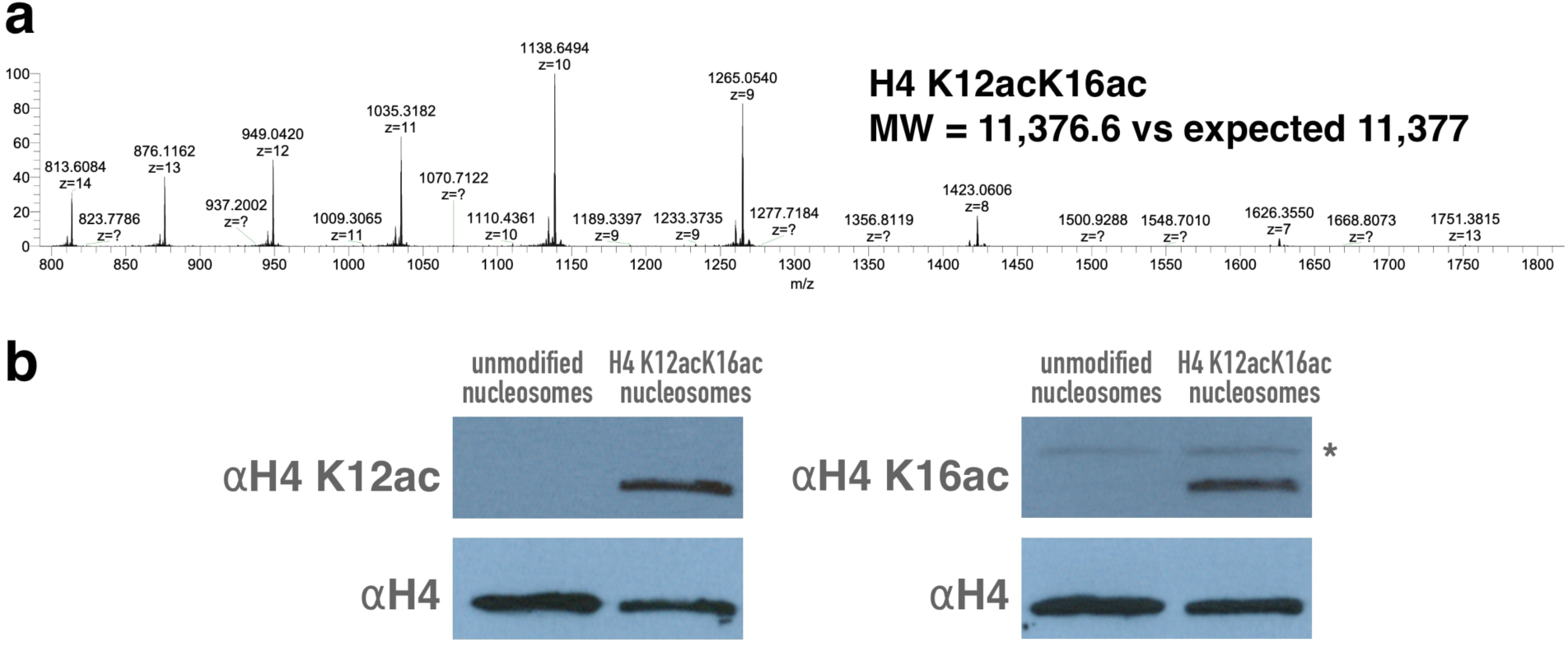
Validation of H4 K12acK16ac nucleosomes. (a) mass spectrometry confirms molecular weight of diacetylated H4 protein, (b) Western blotting using anti-H4 K12ac (Active Motif #61527) and anti-H4 K16ac (Active Motif #39068) antibodies confirm presence of both H4 K12ac and H4 K16ac modifications. The signal identified by the asterisk in the right panel reflects cross-reactivity of the anti-H4 K16ac antibody with another histone protein. The Western blots were reprobed with anti-H4 antibodies (Abcam #ab17036).

**Extended Data Table 1:**
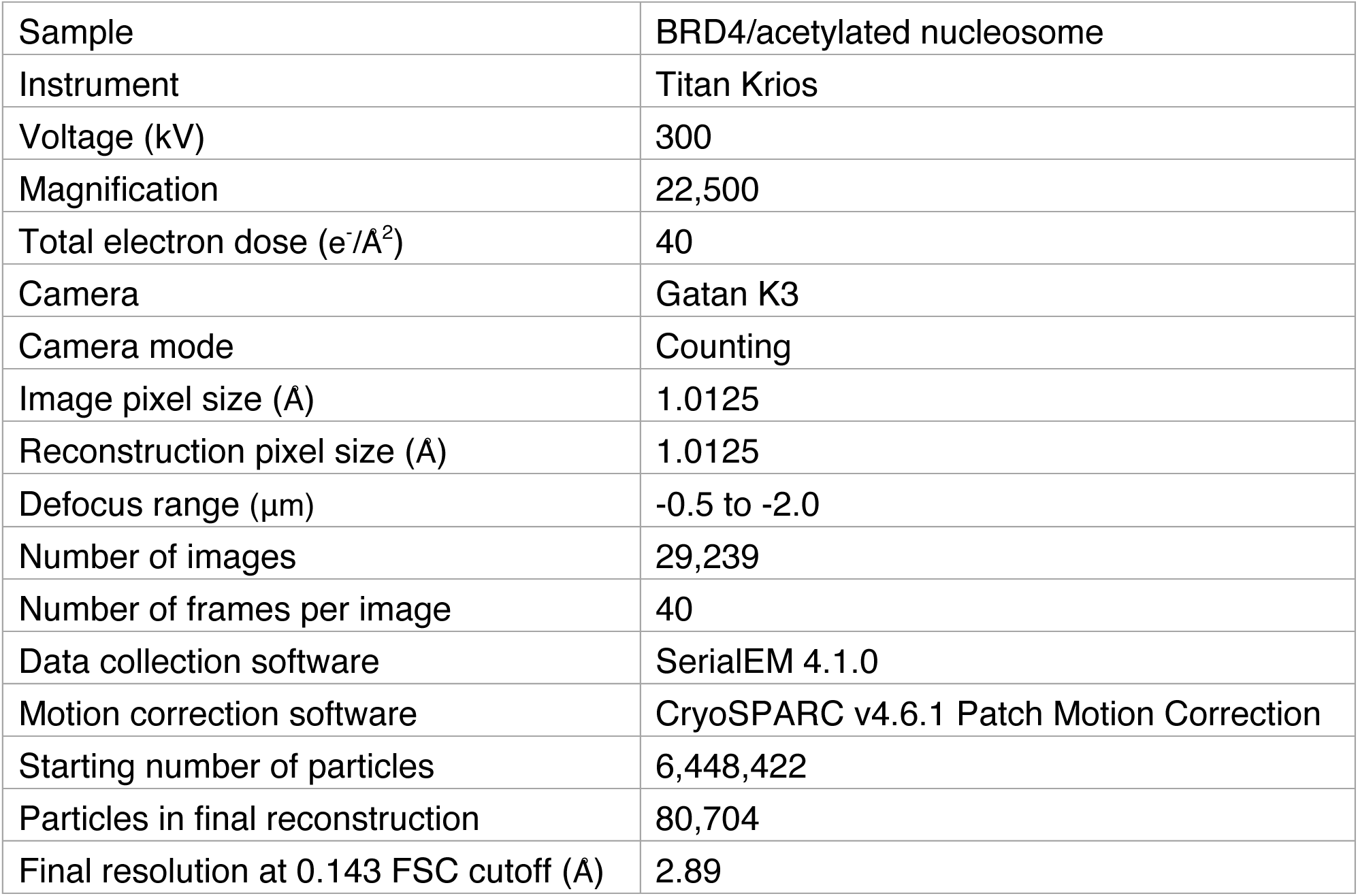
Summary of cryo-EM data collection and refinement.

**Extended Data Table 2:** Model refinement statistics.

| Sample name | BRD4-S/<br>acetylated nucleosome | BRD4-S w basic patch 1/<br>acetylated nucleosome |
| --- | --- | --- |
| Model composition |  |  |
| Atoms | 12,375 | 12,499 |
| Residues: Protein/Nucleotide | 895/255 | 911/255 |
| Model refinement software | Phenix.real_space_refinement |  |
| Model-vs data |  |  |
| CC(mask) | 0.78 | 0.75 |
| CC(box) | 0.84 | 0.84 |
| Resolution estimates (Å) |  |  |
| FSC model (0.143/0.5) Masked | 2.9/3.3 | 3.7/4.5 |
| FSC model (0.143/0.5)<br>Unmasked | 2.9/3.4 | 3.7/6.3 |
| ADP (B-factors, min/max/mean) |  |  |
| Protein | 23.6/177.0/71.6 | 28.4/241.1/112.8 |
| Nucleotide | 82.2/286.3/133.4 | 132.6/545.9/212.4 |
| Bonds (RMSD) |  |  |
| Length (Å) (# > 4σ) | 0.004 | 0.004 |
| Angles (°) (# > 4σ) | 0.758 | 0.749 |
| Scores and Outliers |  |  |
| MolProbity score | 1.06 | 1.13 |
| Clash score | 2.20 | 2.48 |
| Rotamer outliers (%) | 0 | 0 |
| Cβ outliers (%) | 0 | 0 |
| Ramachandran values (%) |  |  |
| Outliers | 0.00 | 0.00 |
| Allowed | 2.29 | 2.48 |
| Favored | 97.71 | 97.52 |

**Extended Data Table 3:**
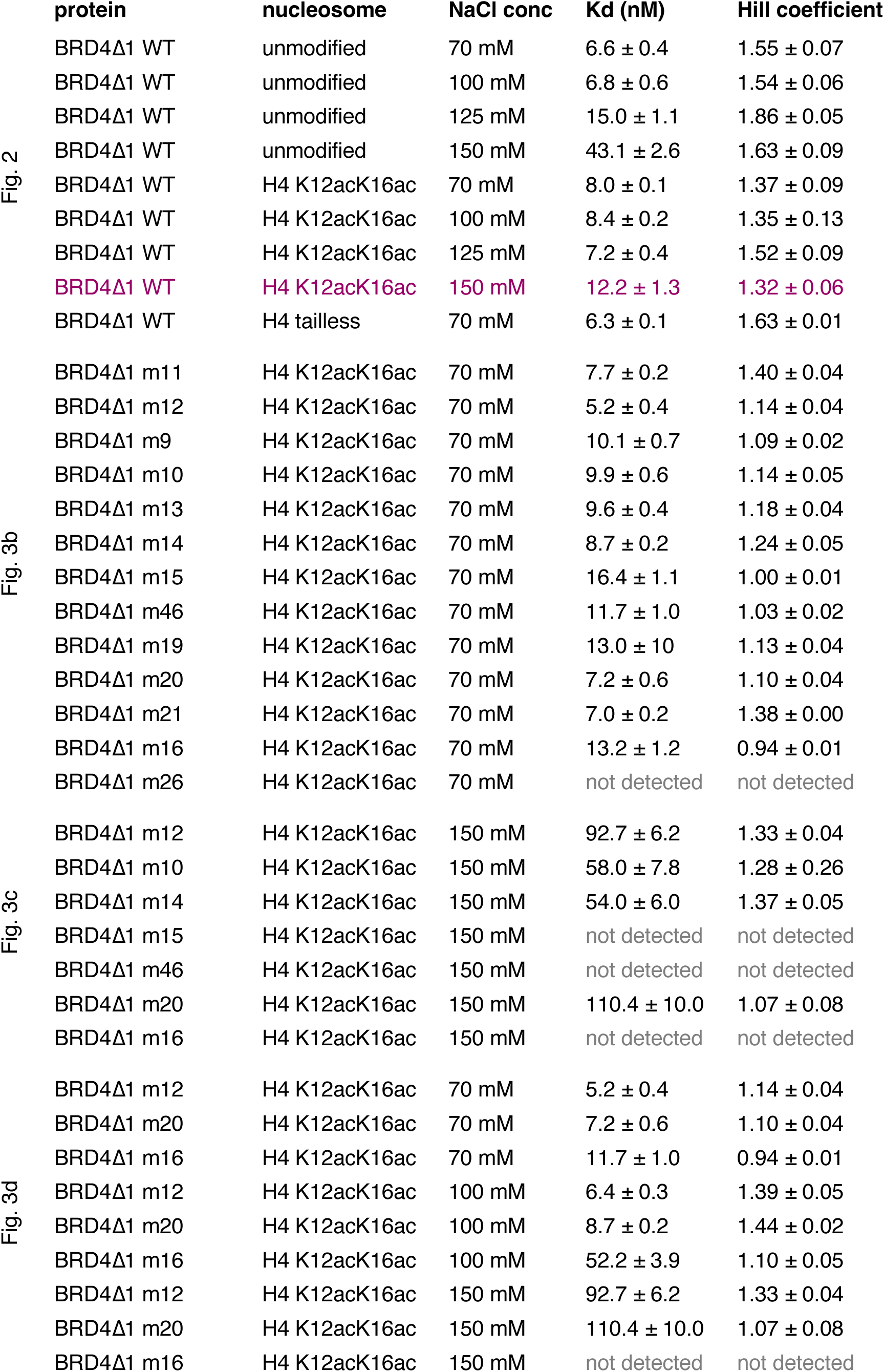
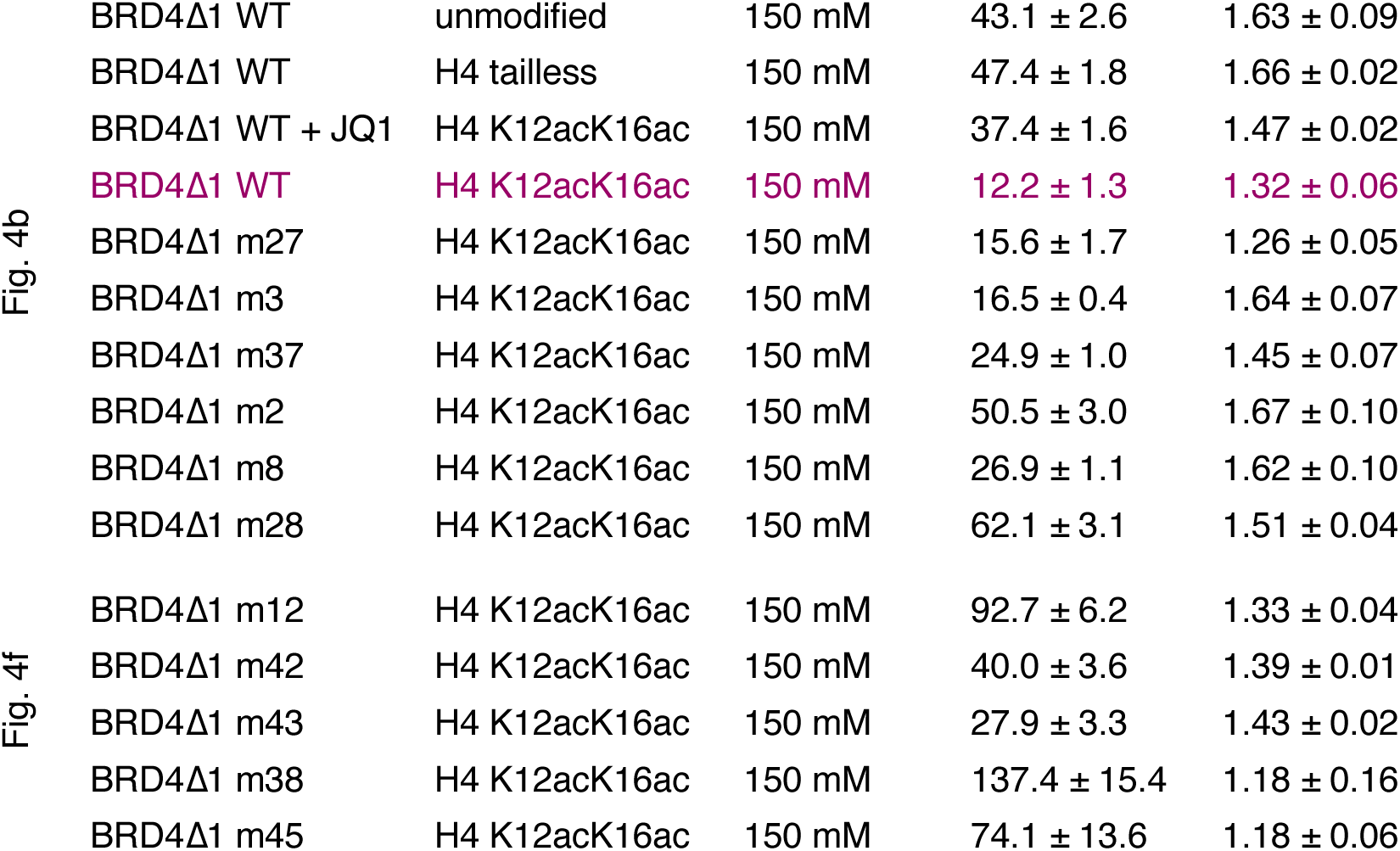
TR-FRET BRD4/nucleosome dissociation constants.

|  | protein | nucleosome | NaCl conc | Kd (nM) | Hill coefficient |
| --- | --- | --- | --- | --- | --- |
| Fig. 2 | BRD4Δ1 WT | unmodified | 70 mM | 6.6 ± 0.4 | 1.55 ± 0.07 |
|  | BRD4Δ1 WT | unmodified | 100 mM | 6.8 ± 0.6 | 1.54 ± 0.06 |
|  | BRD4Δ1 WT | unmodified | 125 mM | 15.0 ± 1.1 | 1.86 ± 0.05 |
|  | BRD4Δ1 WT | unmodified | 150 mM | 43.1 ± 2.6 | 1.63 ± 0.09 |
|  | BRD4Δ1 WT | H4 K12acK16ac | 70 mM | 8.0 ± 0.1 | 1.37 ± 0.09 |
|  | BRD4Δ1 WT | H4 K12acK16ac | 100 mM | 8.4 ± 0.2 | 1.35 ± 0.13 |
|  | BRD4Δ1 WT | H4 K12acK16ac | 125 mM | 7.2 ± 0.4 | 1.52 ± 0.09 |
|  | **BRD4Δ1 WT** | **H4 K12acK16ac** | **150 mM** | **12.2 ± 1.3** | **1.32 ± 0.06** |
|  | BRD4Δ1 WT | H4 tailless | 70 mM | 6.3 ± 0.1 | 1.63 ± 0.01 |
| Fig. 3b | BRD4Δ1 m11 | H4 K12acK16ac | 70 mM | 7.7 ± 0.2 | 1.40 ± 0.04 |
|  | BRD4Δ1 m12 | H4 K12acK16ac | 70 mM | 5.2 ± 0.4 | 1.14 ± 0.04 |
|  | BRD4Δ1 m9 | H4 K12acK16ac | 70 mM | 10.1 ± 0.7 | 1.09 ± 0.02 |
|  | BRD4Δ1 m10 | H4 K12acK16ac | 70 mM | 9.9 ± 0.6 | 1.14 ± 0.05 |
|  | BRD4Δ1 m13 | H4 K12acK16ac | 70 mM | 9.6 ± 0.4 | 1.18 ± 0.04 |
|  | BRD4Δ1 m14 | H4 K12acK16ac | 70 mM | 8.7 ± 0.2 | 1.24 ± 0.05 |
|  | BRD4Δ1 m15 | H4 K12acK16ac | 70 mM | 16.4 ± 1.1 | 1.00 ± 0.01 |
|  | BRD4Δ1 m46 | H4 K12acK16ac | 70 mM | 11.7 ± 1.0 | 1.03 ± 0.02 |
|  | BRD4Δ1 m19 | H4 K12acK16ac | 70 mM | 13.0 ± 10 | 1.13 ± 0.04 |
|  | BRD4Δ1 m20 | H4 K12acK16ac | 70 mM | 7.2 ± 0.6 | 1.10 ± 0.04 |
|  | BRD4Δ1 m21 | H4 K12acK16ac | 70 mM | 7.0 ± 0.2 | 1.38 ± 0.00 |
|  | BRD4Δ1 m16 | H4 K12acK16ac | 70 mM | 13.2 ± 1.2 | 0.94 ± 0.01 |
|  | BRD4Δ1 m26 | H4 K12acK16ac | 70 mM | not detected | not detected |
| Fig. 3c | BRD4Δ1 m12 | H4 K12acK16ac | 150 mM | 92.7 ± 6.2 | 1.33 ± 0.04 |
|  | BRD4Δ1 m10 | H4 K12acK16ac | 150 mM | 58.0 ± 7.8 | 1.28 ± 0.26 |
|  | BRD4Δ1 m14 | H4 K12acK16ac | 150 mM | 54.0 ± 6.0 | 1.37 ± 0.05 |
|  | BRD4Δ1 m15 | H4 K12acK16ac | 150 mM | not detected | not detected |
|  | BRD4Δ1 m46 | H4 K12acK16ac | 150 mM | not detected | not detected |
|  | BRD4Δ1 m20 | H4 K12acK16ac | 150 mM | 110.4 ± 10.0 | 1.07 ± 0.08 |
|  | BRD4Δ1 m16 | H4 K12acK16ac | 150 mM | not detected | not detected |
| Fig. 3d | BRD4Δ1 m12 | H4 K12acK16ac | 70 mM | 5.2 ± 0.4 | 1.14 ± 0.04 |
|  | BRD4Δ1 m20 | H4 K12acK16ac | 70 mM | 7.2 ± 0.6 | 1.10 ± 0.04 |
|  | BRD4Δ1 m16 | H4 K12acK16ac | 70 mM | 11.7 ± 1.0 | 0.94 ± 0.01 |
|  | BRD4Δ1 m12 | H4 K12acK16ac | 100 mM | 6.4 ± 0.3 | 1.39 ± 0.05 |
|  | BRD4Δ1 m20 | H4 K12acK16ac | 100 mM | 8.7 ± 0.2 | 1.44 ± 0.02 |
|  | BRD4Δ1 m16 | H4 K12acK16ac | 100 mM | 52.2 ± 3.9 | 1.10 ± 0.05 |
|  | BRD4Δ1 m12 | H4 K12acK16ac | 150 mM | 92.7 ± 6.2 | 1.33 ± 0.04 |
|  | BRD4Δ1 m20 | H4 K12acK16ac | 150 mM | 110.4 ± 10.0 | 1.07 ± 0.08 |
|  | BRD4Δ1 m16 | H4 K12acK16ac | 150 mM | not detected | not detected |

|  |  |  |  |  |
| --- | --- | --- | --- | --- |
| BRD4Δ1 WT | unmodified | 150 mM | 43.1 ± 2.6 | 1.63 ± 0.09 |
| BRD4Δ1 WT | H4 tailless | 150 mM | 47.4 ± 1.8 | 1.66 ± 0.02 |
| BRD4Δ1 WT + JQ1 | H4 K12acK16ac | 150 mM | 37.4 ± 1.6 | 1.47 ± 0.02 |
| **BRD4Δ1 WT** | **H4 K12acK16ac** | **150 mM** | **12.2 ± 1.3** | **1.32 ± 0.06** |
| BRD4Δ1 m27 | H4 K12acK16ac | 150 mM | 15.6 ± 1.7 | 1.26 ± 0.05 |
| BRD4Δ1 m3 | H4 K12acK16ac | 150 mM | 16.5 ± 0.4 | 1.64 ± 0.07 |
| BRD4Δ1 m37 | H4 K12acK16ac | 150 mM | 24.9 ± 1.0 | 1.45 ± 0.07 |
| BRD4Δ1 m2 | H4 K12acK16ac | 150 mM | 50.5 ± 3.0 | 1.67 ± 0.10 |
| BRD4Δ1 m8 | H4 K12acK16ac | 150 mM | 26.9 ± 1.1 | 1.62 ± 0.10 |
| BRD4Δ1 m28 | H4 K12acK16ac | 150 mM | 62.1 ± 3.1 | 1.51 ± 0.04 |

|  |  |  |  |  |
| --- | --- | --- | --- | --- |
| BRD4Δ1 m12 | H4 K12acK16ac | 150 mM | 92.7 ± 6.2 | 1.33 ± 0.04 |
| BRD4Δ1 m42 | H4 K12acK16ac | 150 mM | 40.0 ± 3.6 | 1.39 ± 0.01 |
| BRD4Δ1 m43 | H4 K12acK16ac | 150 mM | 27.9 ± 3.3 | 1.43 ± 0.02 |
| BRD4Δ1 m38 | H4 K12acK16ac | 150 mM | 137.4 ± 15.4 | 1.18 ± 0.16 |
| BRD4Δ1 m45 | H4 K12acK16ac | 150 mM | 74.1 ± 13.6 | 1.18 ± 0.06 |

## Bibliography

1. Wu, S.-Y. & Chiang, C.-M. The Double Bromodomain-containing Chromatin Adaptor Brd4 and Transcriptional Regulation. Journal of Biological Chemistry 282, 13141– 13145 (2007).

2. Donati, B., Lorenzini, E. & Ciarrocchi, A. BRD4 and Cancer: going beyond transcriptional regulation. Mol. Cancer 17, 164–13 (2018).

3. Cheung, K. L., Kim, C. & Zhou, M.-M. The Functions of BET Proteins in Gene Transcription of Biology and Diseases. Frontiers in Molecular Biosciences 8, (2021).

4. Wang, N., Wu, R., Tang, D. & Kang, R. The BET family in immunity and disease. Sig Transduct Target Ther 6, 1–22 (2021).

5. Altendorfer, E., Mochalova, Y. & Mayer, A. BRD4: a general regulator of transcription elongation. Transcription 13, 70–81 (2022).

6. Eischer, N., Arnold, M. & Mayer, A. Emerging roles of BET proteins in transcription and co-transcriptional processing. WIREs RNA 14, e1734 (2023).

7. Chiang, C.-M. Brd4 engagement from chromatin targeting to transcriptional regulation: selective contact with acetylated histone H3 and H4. F1000 Biol Rep 1, 98 (2009).

8. Sanchez, R., Meslamani, J. & Zhou, M.-M. The bromodomain: From epigenome reader to druggable target. Biochimica et Biophysica Acta (BBA) - Gene Regulatory Mechanisms 1839, 676–685 (2014).

9. Fujisawa, T. & Filippakopoulos, P. Functions of bromodomain-containing proteins and their roles in homeostasis and cancer. Nat Rev Mol Cell Biol 18, 246–262 (2017).

10. Trojer, P. Targeting BET Bromodomains in Cancer. Annual review of cancer biology 6, 313–316 (2022).

11. Ali, H. A. et al. A Comprehensive Review of BET Protein Biochemistry, Physiology, and Pathological Roles. Frontiers in Pharmacology 13, (2022).

12. Rahman, S. et al. The Brd4 Extraterminal Domain Confers Transcription Activation Independent of pTEFb by Recruiting Multiple Proteins, Including NSD3. Molecular and Cellular Biology 31, 2641–2652 (2011).

13. Crowe, B. L. et al. Structure of the Brd4 ET domain bound to a C-terminal motif from γ-retroviral integrases reveals a conserved mechanism of interaction. Proceedings of the National Academy of Sciences 113, 2086–2091 (2016).

14. Zhang, Q. et al. Structural Mechanism of Transcriptional Regulator NSD3 Recognition by the ET Domain of BRD4. Structure 24, 1201–1208 (2016).

15. Konuma, T. et al. Structural Mechanism of the Oxygenase JMJD6 Recognition by the Extraterminal (ET) Domain of BRD4. Sci Rep 7, 16272 (2017).

16. Jang, M. K. et al. The Bromodomain Protein Brd4 Is a Positive Regulatory Component of P-TEFb and Stimulates RNA Polymerase II-Dependent Transcription. Molecular Cell 19, 523–534 (2005).

17. Yang, Z. et al. Recruitment of P-TEFb for Stimulation of Transcriptional Elongation by the Bromodomain Protein Brd4. Molecular Cell 19, 535–545 (2005).

18. Peterlin, B. M. & Price, D. H. Controlling the Elongation Phase of Transcription with P-TEFb. Molecular Cell 23, 297–305 (2006).

19. Winter, G. E. et al. BET Bromodomain Proteins Function as Master Transcription Elongation Factors Independent of CDK9 Recruitment. Molecular Cell 67, 5–18.e19 (2017).

20. Muhar, M. et al. SLAM-seq defines direct gene-regulatory functions of the BRD4-MYC axis. Science 360, 800–805 (2018).

21. Arnold, M., Bressin, A., Jasnovidova, O., Meierhofer, D. & Mayer, A. A BRD4-mediated elongation control point primes transcribing RNA polymerase II for 3′-processing and termination. Molecular Cell 81, 3589–3603.e13 (2021).

22. Zheng, B. et al. Acute perturbation strategies in interrogating RNA polymerase II elongation factor function in gene expression. Genes Dev. 35, 273–285 (2021).

23. Bressin, A. et al. High-sensitive nascent transcript sequencing reveals BRD4-specific control of widespread enhancer and target gene transcription. Nat Commun 14, 4971 (2023).

24. Chiang, C.-M. Nonequivalent Response to Bromodomain-Targeting BET Inhibitors in Oligodendrocyte Cell Fate Decision. Chemistry & Biology 21, 804–806 (2014).

25. Wu, S.-Y. et al. Opposing Functions of BRD4 Isoforms in Breast Cancer. Molecular Cell 78, 1114–1132.e10 (2020).

26. Dey, A., Chitsaz, F., Abbasi, A., Misteli, T. & Ozato, K. The double bromodomain protein Brd4 binds to acetylated chromatin during interphase and mitosis. Proceedings of the National Academy of Sciences 100, 8758–8763 (2003).

27. Wang, R., Li, Q., Helfer, C. M., Jiao, J. & You, J. Bromodomain Protein Brd4 Associated with Acetylated Chromatin Is Important for Maintenance of Higher-order Chromatin Structure. Journal of Biological Chemistry 287, 10738–10752 (2012).

28. Dhalluin, C. et al. Structure and ligand of a histone acetyltransferase bromodomain. Nature 399, 491–496 (1999).

29. Owen, D. J. et al. The structural basis for the recognition of acetylated histone H4 by the bromodomain of histone acetyltransferase Gcn5p. EMBO Journal 19, 6141– 6149 (2000).

30. Morinière, J. et al. Cooperative binding of two acetylation marks on a histone tail by a single bromodomain. Nature 461, 664–668 (2009).

31. Filippakopoulos, P. et al. Selective inhibition of BET bromodomains. Nature 468, 1067–1073 (2010).

32. Filippakopoulos, P. & Knapp, S. Targeting bromodomains: epigenetic readers of lysine acetylation. Nat Rev Drug Discov 13, 337–356 (2014).

33. Ferri, E., Petosa, C. & McKenna, C. E. Bromodomains: Structure, function and pharmacology of inhibition. Biochemical Pharmacology 106, 1–18 (2016).

34. Meslamani, J., Smith, S. G., Sanchez, R. & Zhou, M.-M. Structural features and inhibitors of bromodomains. Drug Discovery Today: Technologies 19, 3–15 (2016).

35. Zaware, N. & Zhou, M.-M. Bromodomain biology and drug discovery. Nat Struct Mol Biol 26, 870–879 (2019).

36. Duan, Y. et al. Targeting Brd4 for cancer therapy: inhibitors and degraders. *Med*. Chem. Commun. 9, 1779–1802 (2018).

37. White, M. E., Fenger, J. M. & Carson, W. E. Emerging roles of and therapeutic strategies targeting BRD4 in cancer. Cellular Immunology 337, 48–53 (2019).

38. Worden, E. J., Hoffmann, N. A., Hicks, C. W. & Wolberger, C. Mechanism of Cross-talk between H2B Ubiquitination and H3 Methylation by Dot1L. Cell 176, 1490–1501.e12 (2019).

39. Chio, U. S. et al. Cryo-EM structure of the human Sirtuin 6–nucleosome complex. Science Advances 9, eadf7586 (2023).

40. Kikuchi, M. et al. Epigenetic mechanisms to propagate histone acetylation by p300/CBP. Nat Commun 14, 4103 (2023).

41. Spangler, C. J. et al. Structural basis of paralog-specific KDM2A/B nucleosome recognition. Nat Chem Biol 19, 624–632 (2023).

42. Jung, M. et al. Affinity Map of Bromodomain Protein 4 (BRD4) Interactions with the Histone H4 Tail and the Small Molecule Inhibitor JQ1. Journal of Biological Chemistry 289, 9304–9319 (2014).

43. Wesley, N. A. et al. Time Resolved-Fluorescence Resonance Energy Transfer platform for quantitative nucleosome binding and footprinting. Protein Science 31, e4339 (2022).

44. Paillisson, A. et al. Bromodomain testis-specific protein is expressed in mouse oocyte and evolves faster than its ubiquitously expressed paralogs *BRD2*, -*3*, and -*4*. Genomics 89, 215–223 (2007).

45. Wu, S.-Y., Lee, A.-Y., Lai, H.-T., Zhang, H. & Chiang, C.-M. Phospho Switch Triggers Brd4 Chromatin Binding and Activator Recruitment for Gene-Specific Targeting. Molecular Cell 49, 843–857 (2013).

46. Filippakopoulos, P. et al. Histone recognition and large-scale structural analysis of the human bromodomain family. Cell 149, 214–231 (2012).

47. Han, X. et al. Roles of the BRD4 short isoform in phase separation and active gene transcription. Nat Struct Mol Biol 27, 333–341 (2020).

48. Larue, R. C. et al. Bimodal high-affinity association of Brd4 with murine leukemia virus integrase and mononucleosomes. Nucleic Acids Research 42, 4868–4881 (2014).

49. McGinty, R. K. & Tan, S. Principles of nucleosome recognition by chromatin factors and enzymes. Curr Opin Struct Biol 71, 16–26 (2021).

50. Makde, R. D., England, J. R., Yennawar, H. P. & Tan, S. Structure of RCC1 chromatin factor bound to the nucleosome core particle. Nature 467, 562–566 (2010).

51. England, J. R., Huang, J., Jennings, M. J., Makde, R. D. & Tan, S. RCC1 uses a conformationally diverse loop region to interact with the nucleosome: a model for the RCC1-nucleosome complex. J Mol Biol 398, 518–529 (2010).

52. McGinty, R. K. & Tan, S. Recognition of the nucleosome by chromatin factors and enzymes. Curr Opin Struct Biol 37, 54–61 (2016).

53. Huth, J. R. et al. The solution structure of an HMG-I(Y)-DNA complex defines a new architectural minor groove binding motif. Nat Struct Biol 4, 657–665 (1997).

54. Miller, T. C. R. et al. A bromodomain–DNA interaction facilitates acetylation-dependent bivalent nucleosome recognition by the BET protein BRDT. Nat Commun 7, 13855 (2016).

55. Baud, M. G. J. et al. A bump-and-hole approach to engineer controlled selectivity of BET bromodomain chemical probes. Science 346, 638–641 (2014).

56. Liu, Z. et al. Discovery, X-ray Crystallography, and Anti-inflammatory Activity of Bromodomain-containing Protein 4 (BRD4) BD1 Inhibitors Targeting a Distinct New Binding Site. J. Med. Chem. 65, 2388–2408 (2022).

57. Armache, K.-J., Garlick, J. D., Canzio, D., Narlikar, G. J. & Kingston, R. E. Structural basis of silencing: Sir3 BAH domain in complex with a nucleosome at 3.0 Å resolution. Science 334, 977–982 (2011).

58. Liu, L. et al. Arginine methylation of BRD4 by PRMT2/4 governs transcription and DNA repair. Science Advances 8, eadd8928 (2022).

59. Liu, Y. et al. Methylation of BRD4 by PRMT1 regulates BRD4 phosphorylation and promotes ovarian cancer invasion. Cell Death Dis 14, 1–14 (2023).

60. Xiong, C. et al. PRMT1-mediated BRD4 arginine methylation and phosphorylation promote partial epithelial–mesenchymal transformation and renal fibrosis. The FASEB Journal 39, e70293 (2025).

61. Sabari, B. R. et al. Coactivator condensation at super-enhancers links phase separation and gene control. Science 361, eaar3958 (2018).

62. Strom, A. R., et al. Interplay of condensation and chromatin binding underlies BRD4 targeting. MBoC 35, ar88 (2024).

63. Zengerle, M., Chan, K.-H. & Ciulli, A. Selective Small Molecule Induced Degradation of the BET Bromodomain Protein BRD4. ACS Chem. Biol. 10, 1770–1777 (2015).

64. You, J., Croyle, J. L., Nishimura, A., Ozato, K. & Howley, P. M. Interaction of the Bovine Papillomavirus E2 Protein with Brd4 Tethers the Viral DNA to Host Mitotic Chromosomes. Cell 117, 349–360 (2004).

65. Tan, S., Kern, R. C. & Selleck, W. The pST44 polycistronic expression system for producing protein complexes in Escherichia coli. Protein Expr Purif 40, 385–395 (2005).

66. Studier, F. W. Protein production by auto-induction in high density shaking cultures. Protein Expr Purif 41, 207–234 (2005).

67. Luger, K., Rechsteiner, T. J. & Richmond, T. J. Expression and purification of recombinant histones and nucleosome reconstitution. Methods Mol Biol 119, 1–16 (1999).

68. Neumann, H. et al. A method for genetically installing site-specific acetylation in recombinant histones defines the effects of H3 K56 acetylation. Molecular Cell 36, 153–163 (2009).

69. Wilkins, B. J. et al. Genetically encoding lysine modifications on histone H4. ACS Chem Biol 10, 939–944 (2015).

70. Kastner, B. et al. GraFix: sample preparation for single-particle electron cryomicroscopy. Nat Methods 5, 53–55 (2008).

71. Grassucci, R. A., Taylor, D. J. & Frank, J. Preparation of macromolecular complexes for cryo-electron microscopy. Nat Protoc 2, 3239–3246 (2007).

72. Punjani, A., Rubinstein, J. L., Fleet, D. J. & Brubaker, M. A. cryoSPARC: algorithms for rapid unsupervised cryo-EM structure determination. Nat Methods 14, 290–296 (2017).

73. Grant, T. & Grigorieff, N. Measuring the optimal exposure for single particle cryo-EM using a 2.6 Å reconstruction of rotavirus VP6. eLife 4, e06980 (2015).

74. Punjani, A., Zhang, H. & Fleet, D. J. Non-uniform refinement: adaptive regularization improves single-particle cryo-EM reconstruction. Nat Methods 17, 1214–1221 (2020).

75. Vasudevan, D., Chua, E. Y. D. & Davey, C. A. Crystal structures of nucleosome core particles containing the ‘601’ strong positioning sequence. J Mol Biol 403, 1–10 (2010).

76. Pettersen, E. F. et al. UCSF ChimeraX: Structure visualization for researchers, educators, and developers. Protein Science 30, 70–82 (2021).

77. Emsley, P., Lohkamp, B., Scott, W. G. & Cowtan, K. Features and development of Coot. Acta Cryst D 66, 486–501 (2010).

78. Afonine, P. V. et al. Real-space refinement in PHENIX for cryo-EM and crystallography. Acta Cryst D 74, 531–544 (2018).

79. Williams, C. J. et al. MolProbity: More and better reference data for improved all-atom structure validation. Protein Science 27, 293–315 (2018).

80. Weiss, J. N. The Hill equation revisited: uses and misuses. The FASEB Journal 11, 835–841 (1997).

81. Virtanen, P. et al. SciPy 1.0: fundamental algorithms for scientific computing in Python. Nat Methods 17, 261–272 (2020).

